# Cytosolic pH controls fungal MAPK signaling and pathogenicity

**DOI:** 10.1101/2022.10.23.513408

**Authors:** Tânia R. Fernandes, Melani Mariscal, Antonio Serrano, David Segorbe, Teresa Fernández-Acero, Humberto Martín, David Turrà, Antonio Di Pietro

## Abstract

In fungi, ambient pH acts as a key regulator of development and virulence. The vascular wilt pathogen *Fusarium oxysporum* uses host alkalinization to promote infection of plant hosts through activation of the invasive growth mitogen-activated protein kinase (MAPK) Fmk1. The molecular events underlying pH-driven MAPK regulation are unknown. Using the ratiometric GFP-based pH sensor pHluorin, we find that both *F. oxysporum* and *Saccharomyces cerevisiae* respond to extracellular alkalinization or acidification with a transitory shift in cytosolic pH (pH_c_) and rapid changes in phosphorylation levels of the three fungal MAPKs Fmk1, Mpk1/Slt2 (cell wall integrity) and Hog1 (hyperosmotic stress). Pharmacological inhibition of the essential plasma membrane H+-ATPase Pma1, which leads to pH_c_ acidification, is sufficient to trigger reprogramming of MAPK phosphorylation even in the absence of an extracellular pH shift. Screening of a subset of *S. cerevisiae* mutants identified the sphingolipid-regulated AGC kinase Ypk1/2 as a key upstream component of pH_c_-modulated MAPK responses. We further show that acidification of pH_c_ in *F. oxysporum* leads to an increase of the long chain base (LCB) sphingolipid dihydrosphingosine (dhSph) and that exogenous addition of dhSph activates Mpk1 phosphorylation. Our results reveal a pivotal role of pH_c_ in the regulation of MAPK signaling and suggest new ways to control fungal growth and pathogenicity.

## Introduction

Ambient pH affects a wide range of biological functions including the acquisition of nutrients and microelements, intracellular signaling and cell growth. As a consequence organisms have evolved intricate mechanisms for sensing and modifying the surrounding pH. Fungi can adapt to fluctuations in pH through conserved signaling cascades such as the alkaline response Pal/Rim signal transduction pathway, which mediates transcriptional regulation by ambient pH and has been extensively studied in the model fungi *Aspergillus nidulans* and *Saccharomyces cerevisiae* (Peñalva *et al.*, 2014). Ambient pH also plays a key role in the control of fungal pathogenicity by modulating the expression of virulence-related genes (Fernandes *et al.*, 2017; Li *et al.*, 2021). Moreover, pathogens have the ability to acidify or alkalinize the host pH during infection by secreting molecules such as organic acids or ammonia, respectively (Alkan *et al.*, 2013; Fernandes *et al.*, 2017). We recently showed that *Fusarium oxysporum*, a soil-borne ascomycete pathogen that causes vascular wilt disease in more than 150 crops (Dean *et al.*, 2012), secretes a functional homologue of the plant regulatory peptide RALF (Rapid Alkalinizing Factor) to trigger host alkalinization and increase its virulence on tomato plants (Masachis *et al.*, 2016; Segorbe *et al.*, 2017). By contrast, acidification of the rhizosphere through secretion of gluconic acid by the bacterial endophyte *Rahnella aquatilis* prevents fungal infection *F. oxysporum* (Palmieri *et al.*, 2020).

Alkalinization was found to promote pathogenicity of *F. oxysporum* by triggering phosphorylation of the conserved mitogen-activated protein kinase (MAPK) Fmk1, which is essential for invasive growth (Masachis *et al.*, 2016; Segorbe *et al.*, 2017). Besides the Fmk1 MAPK cascade, *F. oxysporum* uses the functionally distinct cell wall integrity (CWI) MAPK Mpk1/Slt2 to chemotropically sense plant roots in the soil (Nordzieke *et al.*, 2019; Turrà *et al.*, 2015). How pH controls MAPK signaling during fungal infection is currently unknown. In contrast to ambient pH, which can be subject to dramatic changes, cytosolic pH (pH_c_) is tightly controlled by an elaborate cellular pH homeostatic system (Kane, 2016). In *S. cerevisiae*, pH_c_ acts as a key regulator of cell growth (Dechant *et al.*, 2014), metabolism (Dechant *et al.*, 2010; Peters *et al.*, 2013; Young *et al.*, 2010) and fate (Andrés *et al.*, 2019; Isom *et al.*, 2013). However, the possible link between pH_c_ and MAPK-controlled virulence functions in fungi has not been explored. Here we found that ambient pH controls MAPK signaling via rapid fluctuations of pH_c_. We show that, both in *F. oxysporum* and *S. cerevisiae*, extracellular acidification or pharmacological inhibition of Pma1, the major plasma membrane proton-extruding H^+^-ATPase, triggers a marked decrease in pH_c_ resulting in rapid activation of the CWI MAPK Mpk1. Using conditional yeast mutants, we found that pH_c_-triggered Mpk1/Slt2 activation is dependent on the essential sphingolipid-responsive protein kinase Ypk1, which acts upstream of the CWI MAPK pathway (Levin, 2011). We further show that pH_c_ acidification in *F. oxysporum* leads to an increase in the ceramide long-chain sphingoid base dihydrosphingosine, which in turn induces phosphorylation of Mpk1. These results establish a previously unrecognized role of pH_c_ in the regulation of MAPK signaling and fungal pathogenicity.

## Results

### Ambient pH regulates infection-related development in *F. oxysporum*

Previous work established that an increase of ambient pH promotes infection-related functions in *F. oxysporum* (Masachis *et al.*, 2016). Here we set out to investigate the cellular mechanisms underlying pH-mediated control of fungal pathogenicity. Because alkalinization was previously shown to trigger rapid phosphorylation of the MAPK Fmk1, which is essential for invasive growth and plant infection (Di Pietro *et al.*, 2001; Masachis *et al.*, 2016), we hypothesized that the effect of ambient pH in pathogenicity-related functions could be mediated by changes in MAPK activity. To test this, we first examined the role of the three known *F. oxysporum* MAPKs in invasive hyphal growth across cellophane membranes, a process that correlates directly with fungal pathogenicity on plants (López-Berges *et al.*, 2010). Cellophane penetration assays conducted with the wild type strain and all possible combinations of single and double MAPK mutants (Segorbe *et al.*, 2017) confirmed that, in line with a previous study (Masachis *et al.*, 2016), penetration by the wild type strain is functional at pH 7 but not at pH 5 (**Fig. 1A**). These experiments also confirmed that invasive hyphal growth is strictly dependent on the Fmk1 MAPK, since penetration was abolished in the single and double mutants lacking the *fmk1* gene. We noted that cellophane crossing was also impaired in the *hog1*Δ mutant, which lacks the hyperosmotic stress response MAPK, but restored in the *mpk1*Δ*hog1*Δ double mutant, suggesting that Hog1 contributes to invasive growth whereas Mpk1 has an inhibitory role. In line with this idea, the *mpk1*Δ mutant, in contrast to the wild type and the *mpk1Δ+mpk1* complemented strain, was able to cross the cellophane layer even at the restrictive pH 5 (**Fig. 1A**). Taken together, these results suggest that invasive growth of *F. oxysporum* is controlled by ambient pH and the concerted action of the three MAPKs: Fmk1 is essential while Hog1 and Mpk1 act as positive and negative regulators, respectively, of the invasion process.

**Figure 1.**
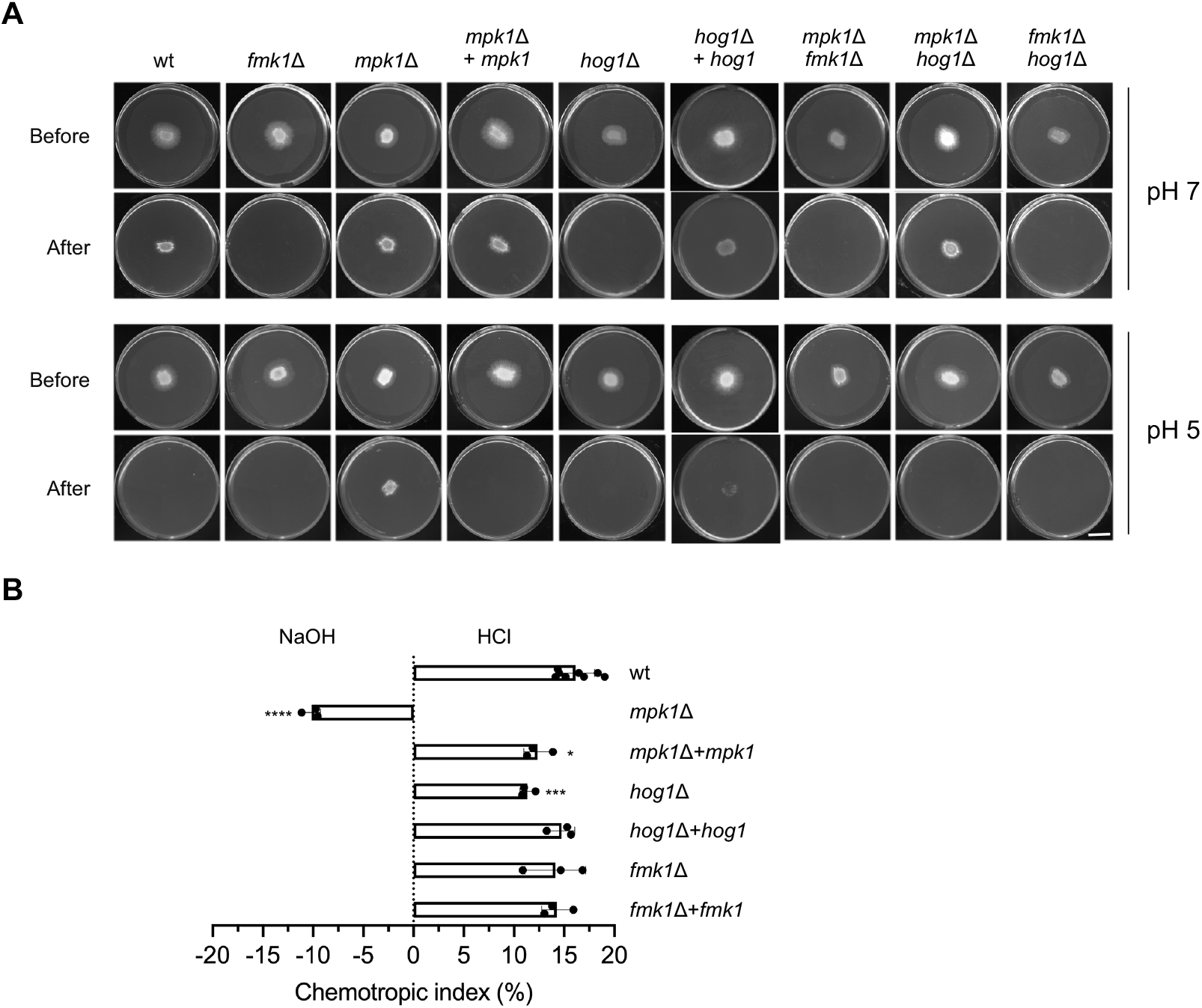
Differential role of MAPK cascades in pH control of fungal infection mechanisms. **A)** Invasive growth of the *F. oxysporum* wild type strain and the indicated single and double MAPK mutants was determined by spot-inoculating the indicated strains on top of a cellophane membrane placed on plates with Potato Dextrose Agar (PDA) buffered at pH 7.0 or 5.0 with 100 mM MES. After 2 days at 28°C plates were imaged (Before), the cellophane with the fungal colony was removed, and plates were incubated for an additional day to visualize the presence of mycelium that had penetrated through the cellophane (After). Images shown are representative of two independent experiments, each with 3 plates per treatment. Scale bar, 2 cm. **B)** Directed growth of germ tubes of the *F. oxysporum* wild type and the indicated mutant strains was determined after 8 h exposure to opposing gradients of 25 mM HCl and NaOH. **** p<0.0001; *** p<0.001; * p<0.05 *versus* wt, according to Welch’s t-test. Data show mean ± s.d from at least three independent experiments (n=500 germ tubes per experiment).

We next examined the possible role of ambient pH in hyphal chemotropism, another infection-related process in *F. oxysporum.* Previous work showed that fungal germ tubes can re-direct growth towards a chemoattractant gradient of peroxidase enzymes released by plant roots, and that this chemotropic response requires the CWI MAPK pathway (Turrà *et al.*, 2015). Here we found that *F. oxysporum* germlings exposed to competing gradients of alkaline and acidic pH grew preferentially towards the acid (**Fig. 1B**). Strikingly, loss of the Mpk1 MAPK led to an inversion of pH tropism, with the *mpk1*Δ mutant growing preferentially towards alkali, while acid tropism was fully restored in the *mpk1*Δ+*mpk1* complemented strain. Thus, *F. oxysporum* hyphae sense pH gradients and re-direct growth towards acidic pH in a Mpk1-dependent manner.

### Shifts in ambient pH trigger rapid re-programming of MAPK phosphorylation

The above findings suggested a possible link between pH and MAPK signaling in the control of infection-related functions. We previously observed rapid phosphorylation of Fmk1 upon extracellular alkalinization (Masachis *et al.*, 2016). Here we found that extracellular acidification had the inverse effect, resulting in dephosphorylation of Fmk1, concomitant with a rapid increase in phosphorylation levels of Mpk1 and Hog1 (**Fig. 2A,B**). A similar response was detected in *S. cerevisiae:* extracellular acidification triggered an increase in phosphorylation of Mpk1 and Hog1 while alkalinization had the opposite effect (**Fig. 2C,D**). To test whether the CWI MAPK Mpk1 contributes to adaptation of *F. oxysporum* to acid stress, we compared growth and survival of the wild type and the *mpk1*Δ mutant under highly acidic conditions. Only minor differences in acid resistance were detected between the wild type and the *mpk1*Δ mutant (**Fig. 2—figure supplement 1**). Taken together these findings indicate that ambient pH fluctuations trigger rapid changes in phosphorylation levels of the three MAPKs, both in *F. oxysporum* and *S. cerevisiae*.

**Figure 2.**
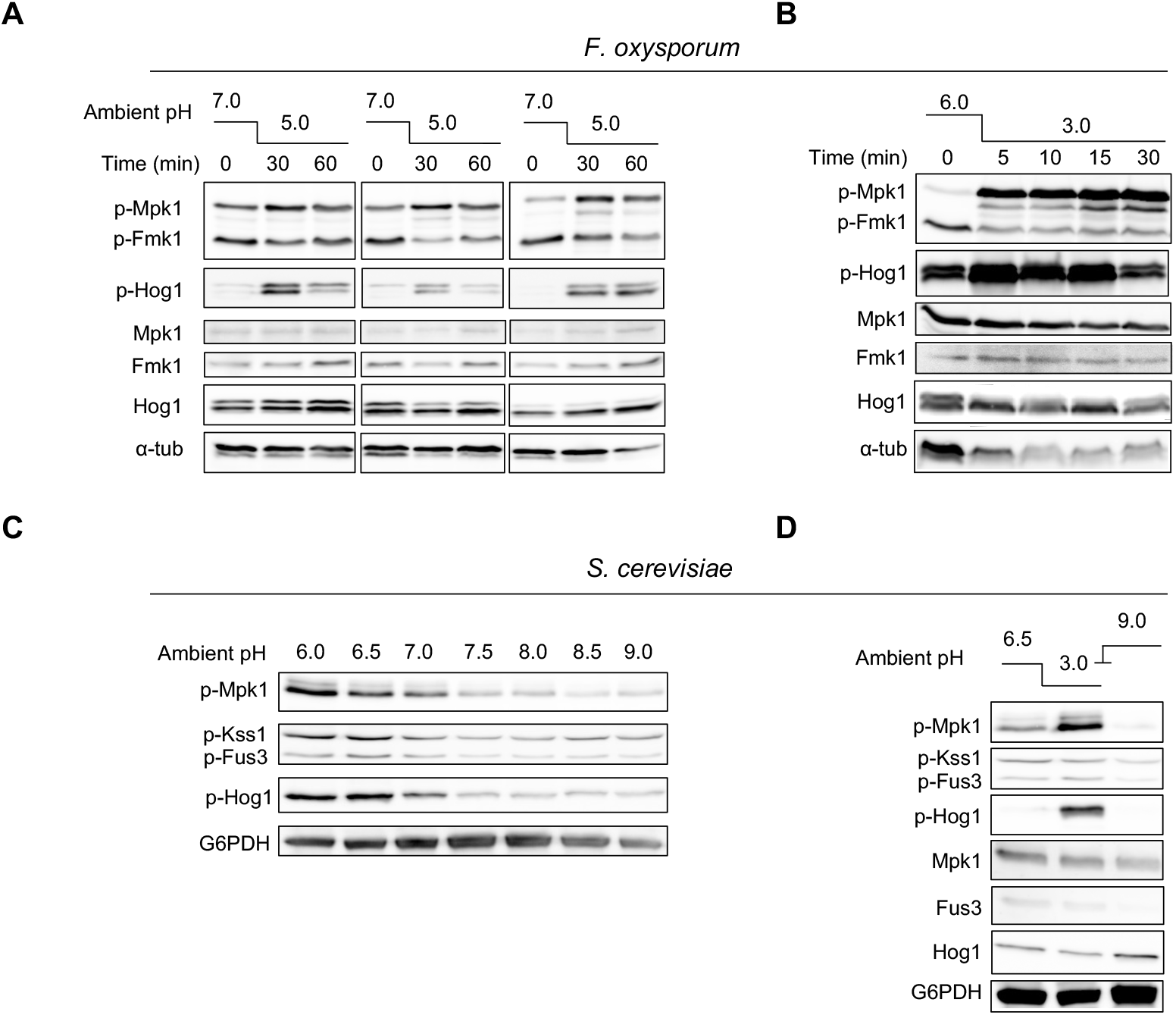
Shifts in ambient pH trigger rapid re-programming of MAPK phosphorylation in *F. oxysporum* and *S. cerevisiae*. **A,B)** *F. oxysporum* microconidia were germinated 15 h at 28°C, either in potato dextrose broth (PDB) buffered at pH 7.0 with 100 mM MES (C); or in yeast extract dextrose medium buffered at pH 7.4 with 20 mM HEPES (YD) and resuspended in KSU buffer at pH 6.0 (D), before the pH of the medium was shifted to 5.0 (C) or 3.0 (D) by adding diluted HCl. Total protein extracts collected at the indicated time points after the pH shift were subjected to immunoblot analysis with anti-phospho-p44/42 or anti-phospho-p38 MAPK antibody to specifically detect phosphorylated p-Mpk1 and p-Fmk1 or p-Hog1, respectively. Anti-Mpk1, anti-Fus3 and anti-Hog1 antibodies were used to detect total MAPK protein levels. Anti-α-tubulin (α-tub) was used as a loading control. Left panel shows immunoblots from 3 independent experiments. **C,D)** *S. cerevisiae* cells grown overnight were either suspended in KSU buffer adjusted to the indicated pH values (A); or suspended in KSU buffer at pH 6.5 and pre-incubated 1 hour at 30°C before shifting the pH of the growth medium to 3.0 or 9.0 by adding diluted HCl or NaOH, respectively (B). Protein extracts were collected 5 min after treatment and subjected to immunoblot analysis with anti-phospho-p44/42 or anti-phospho-p38 MAPK antibodies which specifically detect phosphorylated p-Mpk1 and p-Kss1/p-Fus3 or p-Hog1, respectively. Anti-Mpk1, anti-Fus3 and anti-Hog1 antibodies were used to detect total MAPK protein levels. Anti-G6PDH was used as a loading control.

### pH-triggered changes in MAPK phosphorylation are mediated by rapid fluctuations in cytosolic pH

We next asked how ambient pH controls MAPK activity. We envisaged two possible scenarios that are not mutually exclusive: 1) changes in ambient pH directly or indirectly impinge on pH_c_, which in turn controls MAPK phosphorylation; 2) changes in ambient pH are sensed at the cell surface and directly transduced to the MAPK module.

**Figure 2—figure supplement 1.**
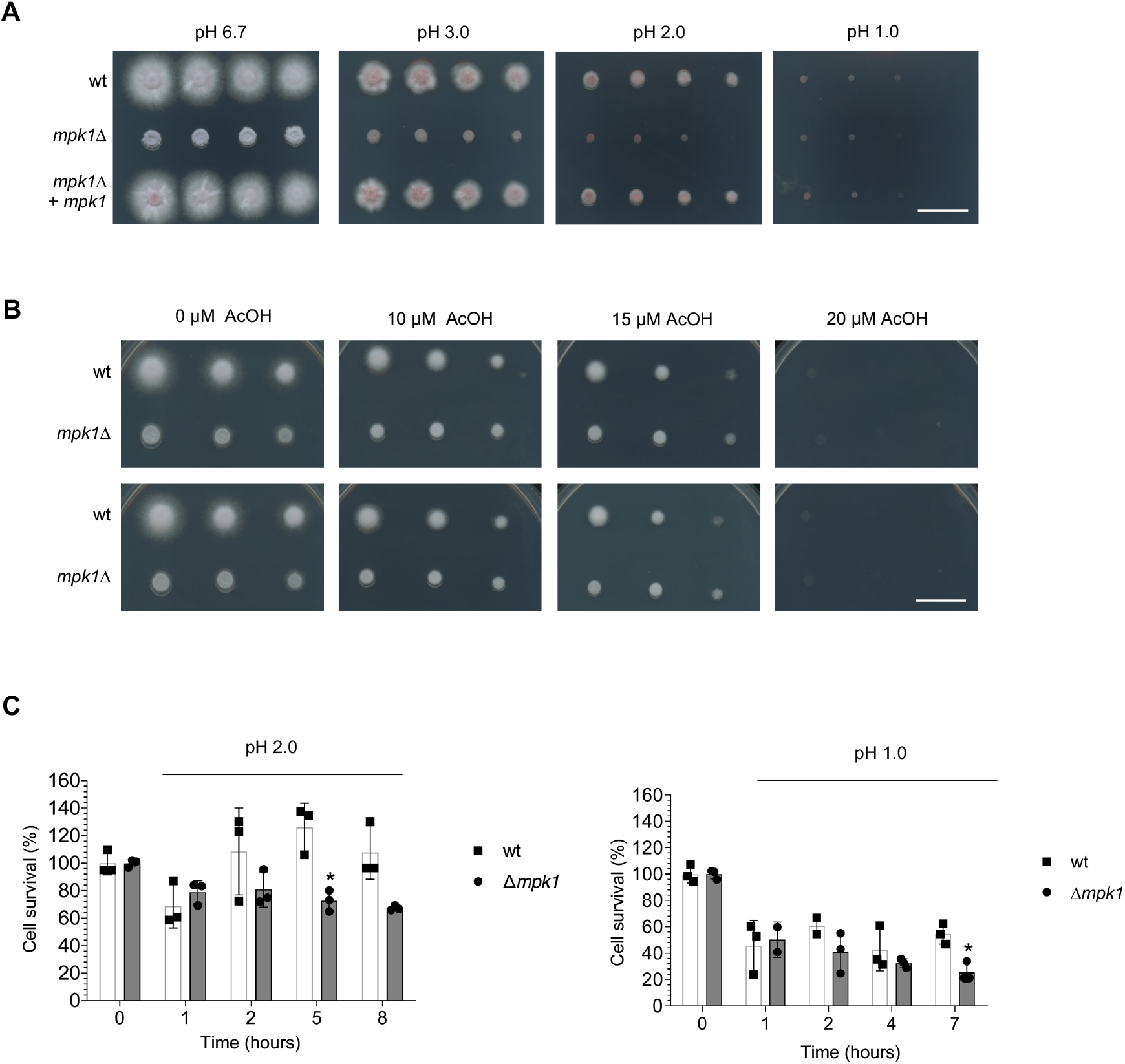
The CWI MAPK Mpk1 has a minor role in adaptation to acidic pH. **A,B)** Serial dilutions of fresh microconidia of the indicated strains were spot-inoculated on PDA plates adjusted to the indicated pH values by adding HCl (A) or supplemented with the indicated concentrations of acetic acid (AcOH) (B). Plates were incubated at 28°C in the dark and imaged after 3 days. Images shown are representative of two independent experiments with three plates each. Scale bar, 2 cm. **C)** The percentage of cell survival of the *F. oxysporum* wild type (wt) and the *mpk1*Δ mutant after the indicated times of exposure to KSU buffer adjusted to pH 2 or 1 by adding HCl was measured by dilution plating and colony counting and normalized to time 0. * p<0.05 *versus* wt according to Welch’s t-test. Data show the mean ± s.d. of three replicate microwells.

To follow pH_c_ in real time, we generated *F. oxysporum* strains expressing the genetically encoded pH-sensor pHluorin, a GFP-derivative that allows *in vivo* ratiometric measurement of pH_c_, driven by the strong constitutive *A. nidulans gpdA* promoter. A pHluorin-expressing transformant displaying high fluorescence levels (**Fig. 3A**) was subjected to *in vivo* calibration with buffers at different pH values, confirming the pH sensitivity and spectral characteristics of the expressed pHluorin protein (**Fig. 3B**). Independent measurements by confocal microscopy and spectrofluorometry in 96-well microtiter plates revealed a uniform distribution of pHluorin in the cytosol of *F. oxysporum* germlings, with a pH_c_ of 7.3 in our standard experimental conditions (**Fig. 3A,C**).

**Figure 3.**
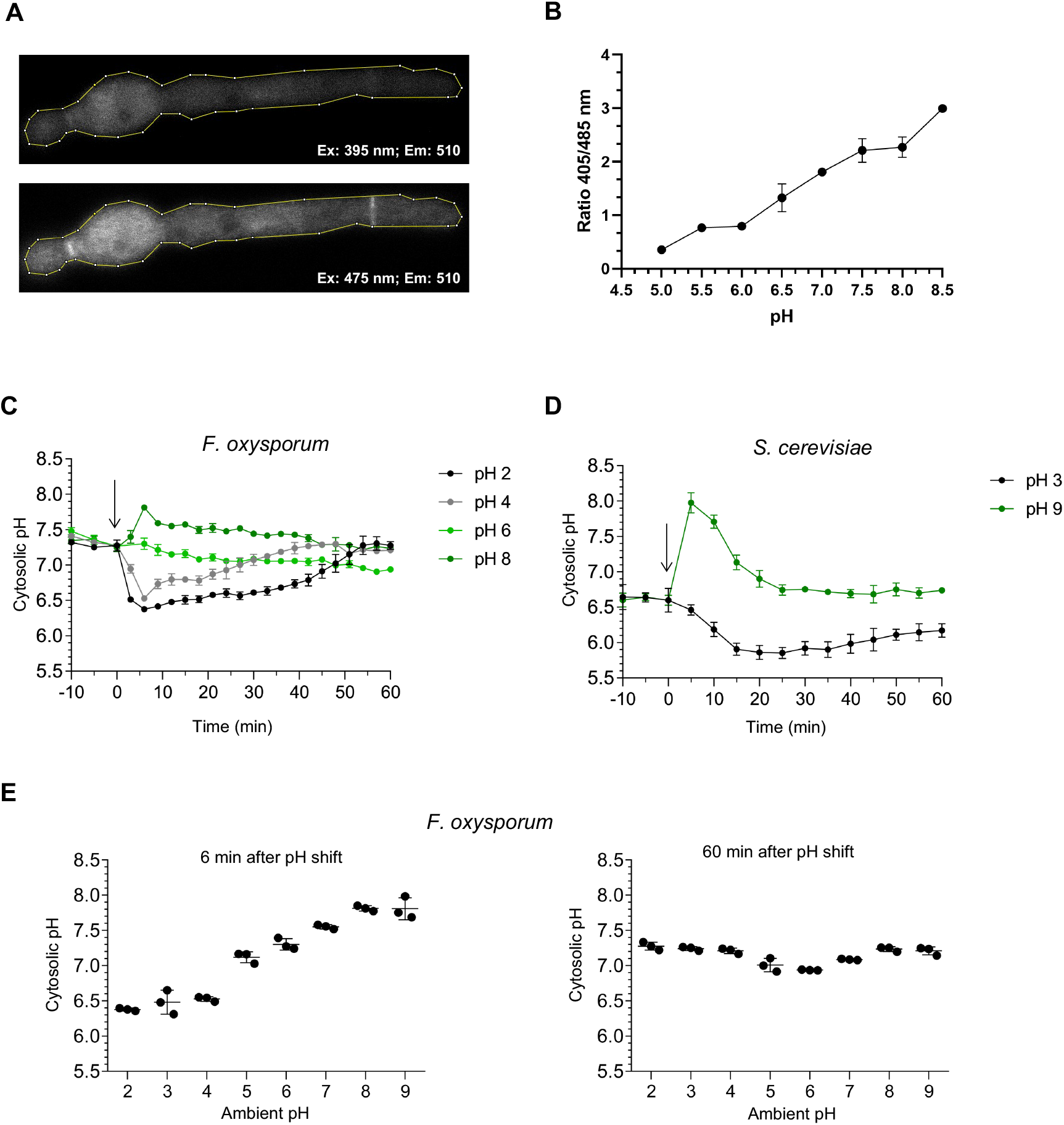
A shift in ambient pH triggers transient changes in cytosolic pH. **A,B)** *F. oxysporum* microconidia germinated 15 hours at 28°C in YD medium buffered at pH 7.4 with 20 mM HEPES or overnight in YPD medium were subjected to analysis of cytosolic pH (pH_c_) using a Zeiss LSM880 laser confocal microscope equipped with diode (405 nm) and Argon (488 nm) lasers, using a Plan Apo 63x oil 1.4 NA objective. A line delimiting the shape of the hypha was drawn (A), fluorescence intensity within the line was measured at 405 and 488 nm wavelength, and the 405/488 nm ratio was calculated for each pH value of the calibration curve (B). **C,D)** The effect of ambient pH shifts on pH_c_ was measured in *F. oxysporum* (A) and *S. cerevisiae* (B) strains expressing the ratiometric pH probe pHluorin. *F. oxysporum* microconidia (A) or *S. cerevisiae* cells (B) were grown 15 h in YD or in Yeast peptone dextrose (YPD) medium, respectivly, resuspended in KSU buffer at pH 6.0 or 6.5, respectively, transferred to microwells and pre-incubated 50 min at 28°C or 30°C, respectively, before shifting the pH of the medium to the indicated values as described in Fig. 2. pH_c_ was monitored spectrofluorometrically every 3 minutes starting 10 minutes before the pH shift. The ratio between the emission intensities at 510 nm after excitation at 395 nm and 475 nm was calculated and normalized to the standard curve (see Suppl. Fig. 2). Data show the mean ± s.d of three replicate microwells from one representative experiment. Experiments were performed at least twice with similar results. **E)** *F. oxysporum* microconidia were pretreated as described in (A) and pH_cyt_ was measured spectrofluorometrically 6 and 60 minutes after shifting the pH of the medium to the indicated values. Data show the mean ± s.d. of three replicate microwells from one representative experiment. Experiments were performed twice with similar results.

We also generated transformants of *S. cerevisiae* strain BY4741 with a plasmid containing the *pHluorin2* gene driven by the *TEF1* promotor (Isom *et al.*, 2013). Spectrofluorometric measurements of these transformants revealed a pH_c_ of around 6.6 (**Fig. 3D**), which is similar to that reported in previous studies (Isom *et al.*, 2013).

We next asked whether pH_c_ is affected by changes in ambient pH. Acidification or alkalinization of the external medium triggered a marked down- or upshift, respectively, of pH_c_ both in *F. oxysporum* and *S. cerevisiae* (**Fig. 3C,D**). The most extreme down- or upshifts of external pH tested in *F. oxysporum* (from pH 6.0 to pH 2.0 or pH 9.0) led a fall or rise, respectively, of approximately one or 0.5 units in pH_c_, from 7.3 to 6.38 ± 0.02 or to 7.81 ± 0.16, respectively (**Fig. 3E**). The fluctuations in pH_c_ were rapid and transient, with a maximum amplitude around 6 minutes after the change in ambient pH and a gradual return to the homeostatic value of 7.3 (**Fig. 3C-E**). These findings suggest the existence of a robust pH homeostasis control which protects the fungal cell during prolonged exposure to extreme ambient pH values.

How do shifts in ambient pH cause changes in pH_c_? Previous work identified the plasma membrane H^+^-ATPase Pma1 as a master regulator of pH_c_ homeostasis in fungi (Kane, 2016). Here we found that a downshift of external pH from 7.0 to 5.0 led to a rapid decrease of Pma1 H^+^-ATPase activity (**Fig. 4A**), suggesting a possible implication of Pma1 the intracellular acidification observed in **Fig. 3C**. Furthermore, pharmacological inhibition of Pma1 H^+^-ATPase activity with the specific inhibitor diethylstilbestrol (DES) (Kahm *et al.*, 2012; Moskvina *et al.*, 1999) caused a rapid (5 min) and sustained drop in pH_c_ of approximately one pH unit, both in *F. oxysporum* (**Fig. 4B,C; Fig. 4—figure supplement 1A**) and in *S. cerevisiae* (**Fig. 4D**). DES-induced pH_c_ acidification was independently confirmed by confocal microscopy measurements (**Fig. 4—figure supplement 1B**).

**Figure 4.**
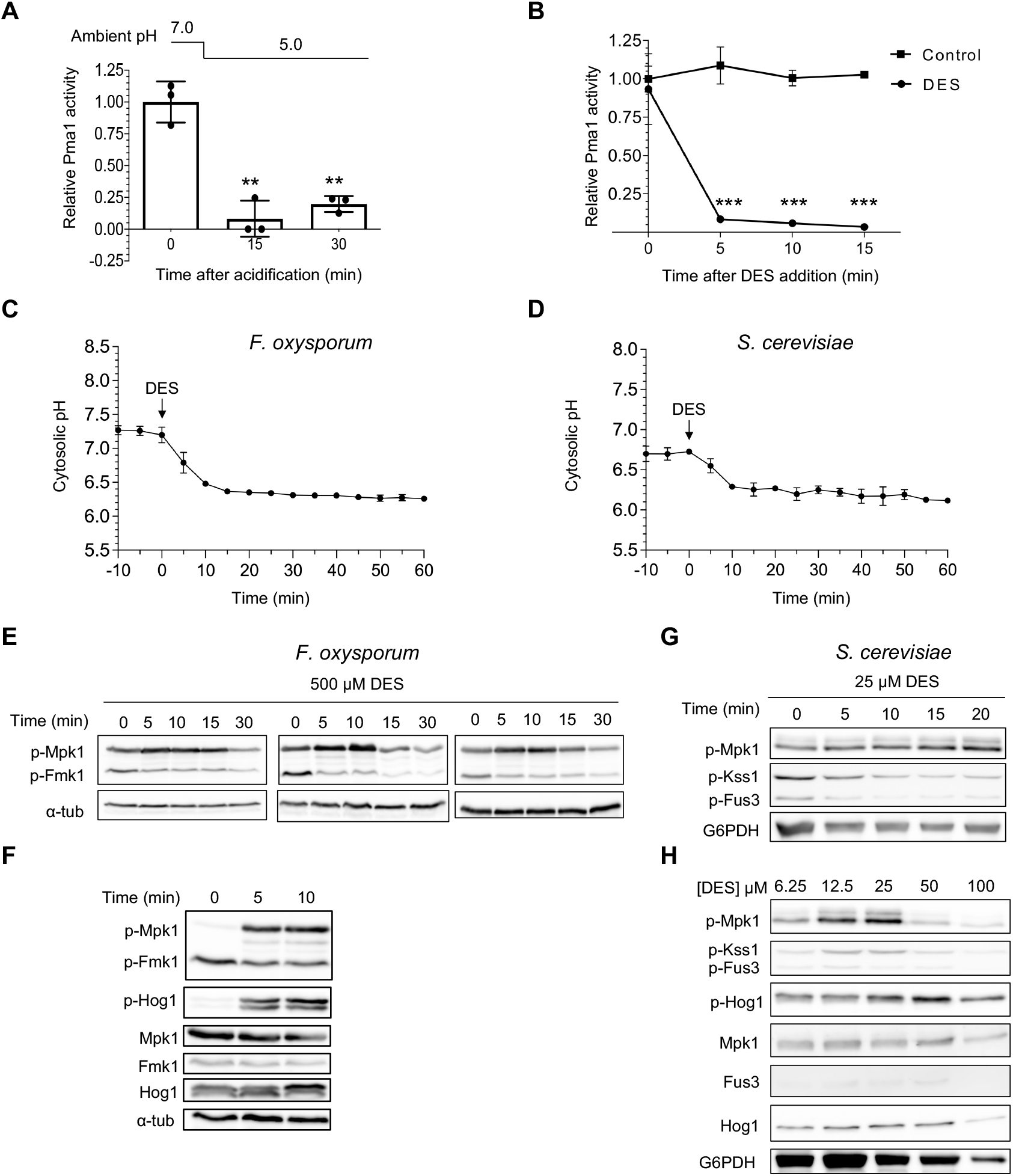
Cytosolic pH controls MAPK phosphorylation. **A,B)** Activity of the major *F. oxysporum* plasma membrane H^+^-ATPase Pma1 is inhibited by acidic ambient pH and by the specific inhibitor diethylstilbestrol (DES). (A,B) Microconidia of *F. oxysporum* were germinated as described in Fig. 2A before shifting the pH of the medium from 7 to 5 with diluted HCl (A) or adding 500 μM DES (B). Total membrane fraction was isolated from mycelia harvested at the indicated time points and H^+^-ATPase activity of Pma1 was measured and normalized to time 0. *** p<0.001; ** p<0.01 according to Welch’s t-test *versus* time 0 (A) or untreated control (B). Data show the mean ± s.d. of three biological replicates from one representative experiment. Experiments were performed twice with similar results. **C,D)** Pma1 inhibition by DES triggers rapid and sustained acidification of pH_c_. *F. oxysporum* microconidia (C) or *S. cerevisiae* cells (D) were pretreated as described in Fig. 3 before adding 500 or 25 μM DES, respectively. pH_c_ was monitored spectrofluorometrically every 5 minutes starting 10 minutes before DES addition. Data show the mean ± s.d. of three independent replicates from one representative experiment. Experiments were performed three times with similar results. **E-H)** DES-triggered pH_c_ acidification leads to rapid Mpk1 phosphorylation and Fmk1 dephosphorylation. *F. oxysporum* microconidia (E,F) or *S. cerevisiae* cells (G,H) were subjected to DES treatment as described in (C,D). Total protein extracts collected at the indicated times after addition of the indicated concentrations of DES were analyzed by immunoblot with different antibodies as indicated in Fig. 2. (E) shows immunoblots from 3 independent biological experiments.

Pharmacological inhibition of Pma1 provides a powerful tool to manipulate pH_c_ independently of changes in extracellular pH. We next tested the effect of DES-triggered intracellular acidification on MAPK phosphorylation and detected a rapid increase in Mpk1 phosphorylation concomitant with dephosphorylation of Fmk1 (**Fig. 4E,F; Fig. 4—figure supplement 1G**). This response mimics that observed previously for extracellular acidification (see **Fig. 2A**). We further noted that treatment of *F. oxysporum* with the proton ionophore carbonyl cyanide-p-trifluoromethoxylphenylhydrazone (FCCP), which results in intracellular acidification, also induced rapid phosphorylation of Mpk1 and dephosphorylation of Fmk1 (**Fig. 4—figure supplement 1C,D**). In *S. cerevisiae*, treatment with DES also triggered rapid phosphorylation of Mpk1 and dephosphorylation of the Fmk1 orthologs Kss1 and Fus3 (**Fig. 4G,H**). We conclude that MAPK phosphorylation is regulated by pH_c_ and that this mechanism is conserved in fungi.

**Figure 4—figure supplement 1.**
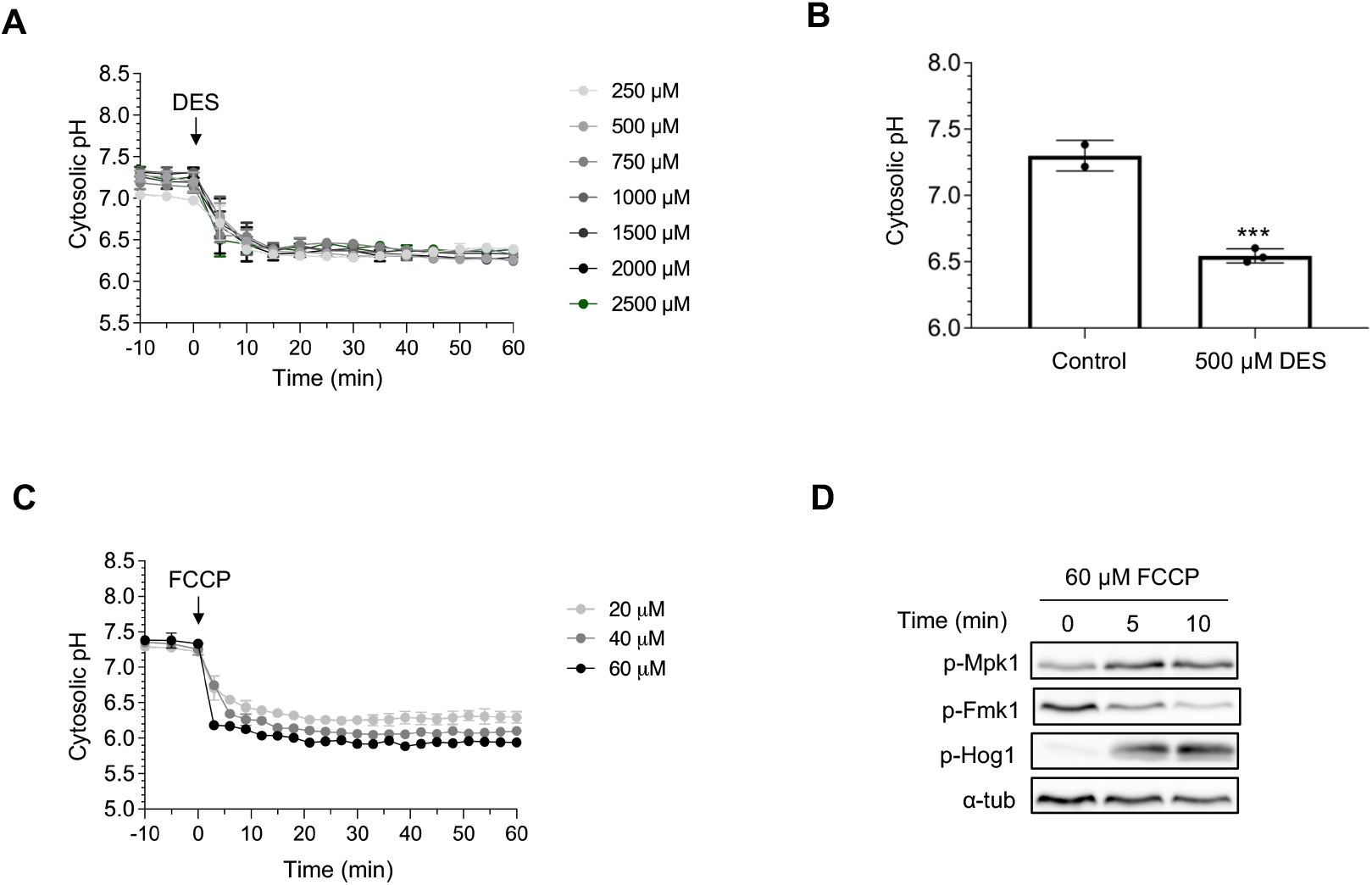
Pma1 inhibition by DES causes rapid acidification of pH_c_ and cell death. **A,B)** Pma1 inhibition by DES causes rapid and sustained acidification of pH_c_. *F. oxysporum* microconidia were pretreated as described in Fig. 3 before adding the indicated concentrations of DES to the medium. pH_c_ was monitored spectrofluorometrically (A) or by confocal microscopy (B) as described in Fig. 3 (C) or (B), respectively. Data show the mean ± s.d. of three independent replicate microwells from one representative experiment. Experiments were performed twice with similar results. **C)** Membrane depolarization by carbonyl cyanide p-trifluoromethoxyphenylhydrazone (FCCP) causes rapid and sustained acidification of pH_c_. *F. oxysporum* microconidia were pretreated as described in Fig. 3 before adding the indicated concentrations of FCCP to the medium. pH_c_ was monitored spectrofluorometrically as described in Fig. 3. Data show the mean ± s.d. of three independent replicate microwells from one representative experiment. Experiments were performed twice with similar results. **D)** *F. oxysporum* microconidia were pretreated as described in Fig. 3, and 60 μM FCCP was added to the medium. Total protein extracts collected at the indicated times were subjected to immunoblot analysis with different antibodies as indicated in Fig. 2.

### The Pal/Rim pathway partially contributes to acid-triggered Mpk1 activation

To ask how pH_c_ controls MAPK activity, we first examined the Pal/Rim pathway, a broadly conserved mechanism of ambient pH sensing and response in fungi. Upon a shift to alkaline pH, the seven transmembrane domain receptor PalH/Rim21 initiates a signaling cascade resulting in proteolytic activation of the zinc finger transcription factor PacC/Rim101, that acts both as an activator of alkaline-expressed and a repressor of acidic-expressed genes (Peñalva *et al.*, 2014; Peñalva *et al.*, 2008). To test the role of the Pal/Rim pathway in pH-mediated MAPK signaling of *F. oxysporum*, we generated *pacC*Δ and *palH*Δ deletion mutants both in the wild type and the pHluorin-expressing backgrounds (**Fig. 5—figure supplement 1A-D**). In line with previous reports (Caracuel *et al.*, 2003; Peñalva *et al.*, 2008), the *pacC*Δ and *palH*Δ mutants exhibited a severe growth defect at high pH but were unaffected in virulence on tomato plants (**Fig. 5A,B)**. Moreover, pH_c_ dynamics of the *pacC*Δ and*palH*Δ mutants in response to external acidification or DES treatment was similar to that of the wild type (**Fig. 5C-F)**, although acid-induced Mpk1 phosphorylation was somewhat delayed in the *pacC*Δ and *palH*Δ mutants compared to the wild type strain (**Fig. 5G)**. This finding suggests that the Pal/Rim pathway could play a minor role in acid pH-mediated MAPK regulation.

**Figure 5.**
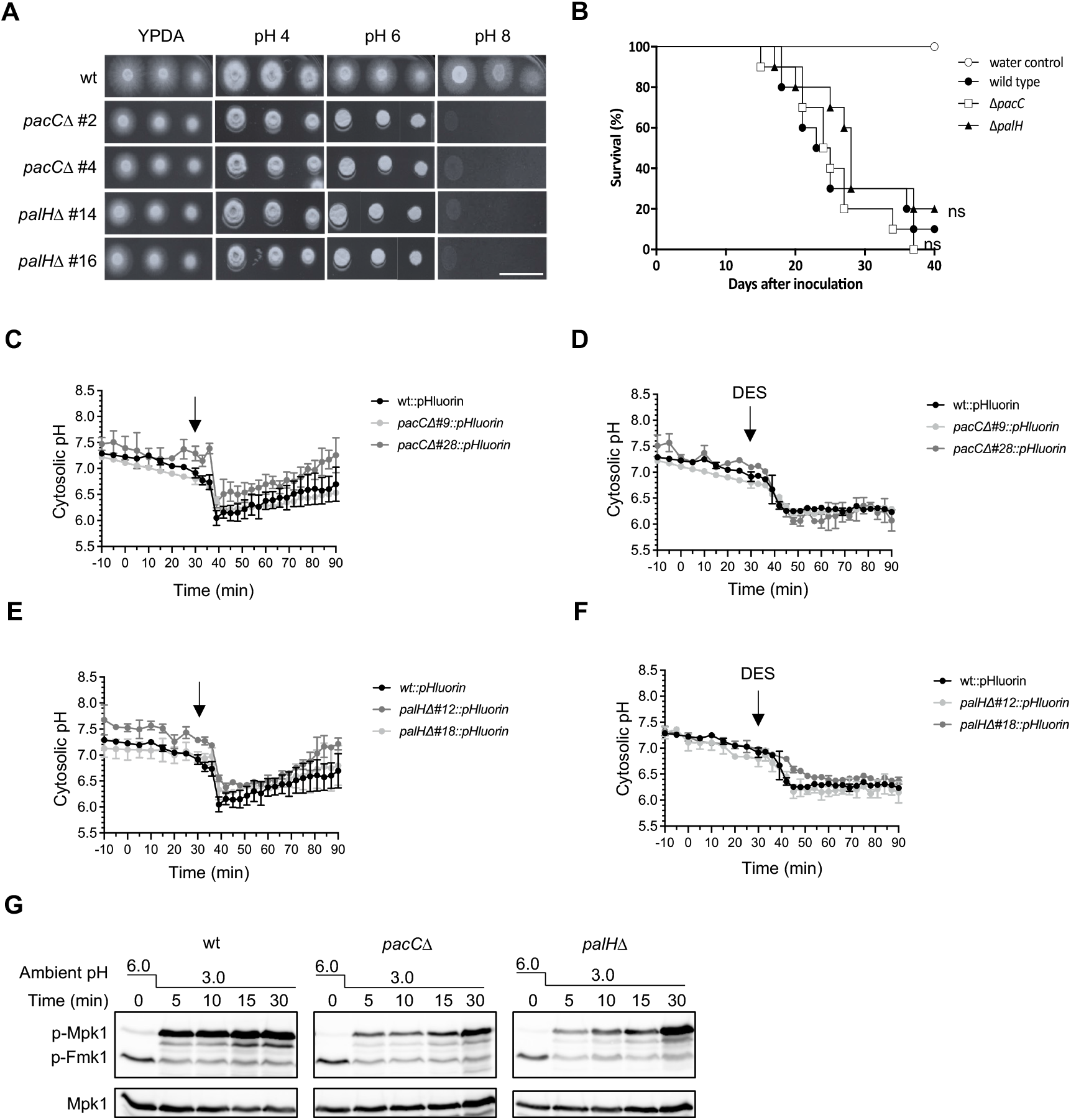
The Pal/PacC pathway is required for adaptation to high pH and contributes to acidification-triggered Mpk1 activation. **A)** Serial dilutions of fresh microconidia of the indicated strains were spot-inoculated on plates containing YPDA medium buffered to the indicated pH with citrate-phosphate buffer. Plates were incubated at 28°C in the dark and imaged after 2 days. Images shown are representative of two independent experiments with three plates each. Scale bar, 2 cm. **B)** Kaplan-Meier plot showing the survival of tomato plants inoculated with the wild type strain or the indicated mutants. Groups of 10 plants were used. Data shown are from one representative experiment. Experiments were performed twice with similar results. ns = non-significant *versus* wild type strain, according to log-rank test. **C-F)** Microconidia of the indicated *F. oxysporum* strains were pretreated as described in Fig. 2 before shifting the pH of the medium from 6.0 to 3.0 by adding diluted HCl (A,C) or adding 500 μM DES (B,D). pH_cy_t was monitored spectrophluorometrically every 3 minutes. Data show the mean ± s.d. of three independent replicates from one representative experiment. Experiments were performed twice with similar results. **G)** The indicated *F. oxysporum* strains were subjected to acidification of ambient pH as described in (A). Total protein extracts collected at the indicated times after the pH shift were analyzed byo immunoblot with anti-phospho-p44/42 to specifically detect phosphorylated p-Mpk1 and p-Fmk1. Anti-α-tubulin (α-tub) was used as a loading control.

**Figure 5—figure supplement 1.**
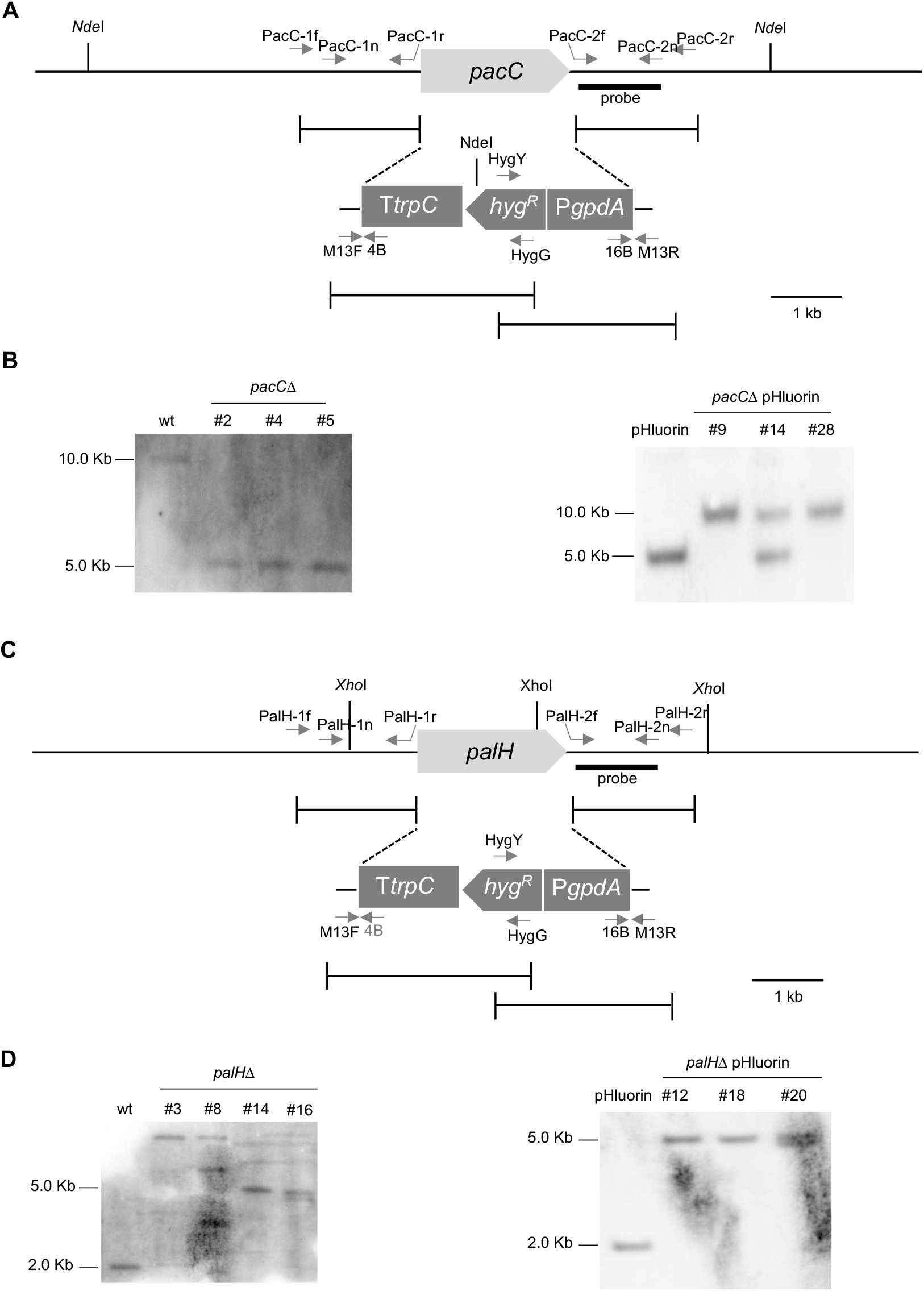
Targeted deletion of *pacC* and *palH* in *F. oxysporum.* **A,C)** Schematic diagram showing targeted deletion of the *F. oxysporum pacC* (A) and *palH* (C) genes using the split-marker method. Gene knockout constructs were obtained by fusion PCR. Relative positions of restriction sites and Southern probes as well as of the PCR primers used are indicated. *hygR*, hygromycin resistance gene; *PgpdA, gpdA* promoter; *TtrpC, trpC* terminator (both from *A. nidulans).* **B,D)** Genomic DNA of independent transformants obtained in the wild type (wt, left panels) or the pHluorin-expressing background (right panels) was treated with *NdeI* (B) or *Xho*I (D), separated on 0.7% agarose gels, transferred to nylon membranes and hybridized with DIG labelled DNA probes from the indicated genes. Molecular weights of the hybridizing bands are indicated on the left.

### The Pkh-Ypk upstream branch is required for acid pH_c_-triggered Mpk1 activation

To gain insights into the mechanisms operating upstream of Mpk1, we used western blot analysis of DES-treated cells to screen a collection of *S. cerevisiae* mutants in known components of the CWI pathway (**Fig. 6A**) for defects in acid pH_c_-triggered MAPK responses. Deletion mutants in the cell surface sensors Wsc1, Mid2 or Mtl1, the downstream guanine exchange factor Rom2 or a *rho1* temperature sensitive (ts) mutant were largely unaffected in DES-triggered Mpk1 phosphorylation and Fus3/Kss1 dephosphorylation (**Fig. 6B**). By contrast, mutants lacking the MAPKKK Bck1 or carrying a temperature sensitive allele of Pkc1 had constitutively low phosphorylation levels of Mpk1, although the *bck1*Δ mutant still exhibited a detectable dephosphorylation response of Fus3/Kss1. We next examined the role of the Pkh-Ypk upstream branch in DES-triggered MAPK regulation. In *S. cerevisiae* the two AGC kinase paralogs Ypk1/2 are phosphorylated by the 3-phosphoinositide-dependent kinase 1 paralogs Pkh1/Pkh2 and the target of rapamycin complex 2 (TORC2) (Niles *et al.*, 2012; Niles and Powers, 2014) (see **Fig. 6A**). Here we found that single deletion mutants in *PKH1, PKH2, YPK1* or *YPK2* genes were largely unaffected in DES-triggered Mpk1 phosphorylation and Kss1/Fus3 dephosphorylation, possibly due to functional redundancy of these gene paralogs (**Fig. 6C**). By contrast, a mutant carrying a deletion of *ypk2* and a temperature sensitive *ypk1* allele *(ypk2Δ ypk1-ts)* was unable to activate Mpk1 in response to DES-triggered intracellular acidification when shifted to the restrictive temperature previous to DES addition (**Fig. 6B**). Likewise, a *ypk2*Δ mutant expressing the analog-sensitive (AS) *ypk1^L424G^* allele (Berchtold *et al.*, 2012) was defective in DES-triggered Mpk1 phosphorylation when treated with the PP1 analog 1-NM-PP1 (**Fig. 6D,E**). Together, these results suggest that acid pH_c_-triggered activation of Mpk1 is mediated by the Pkh-Ypk upstream branch of the CWI cascade.

**Figure 6.**
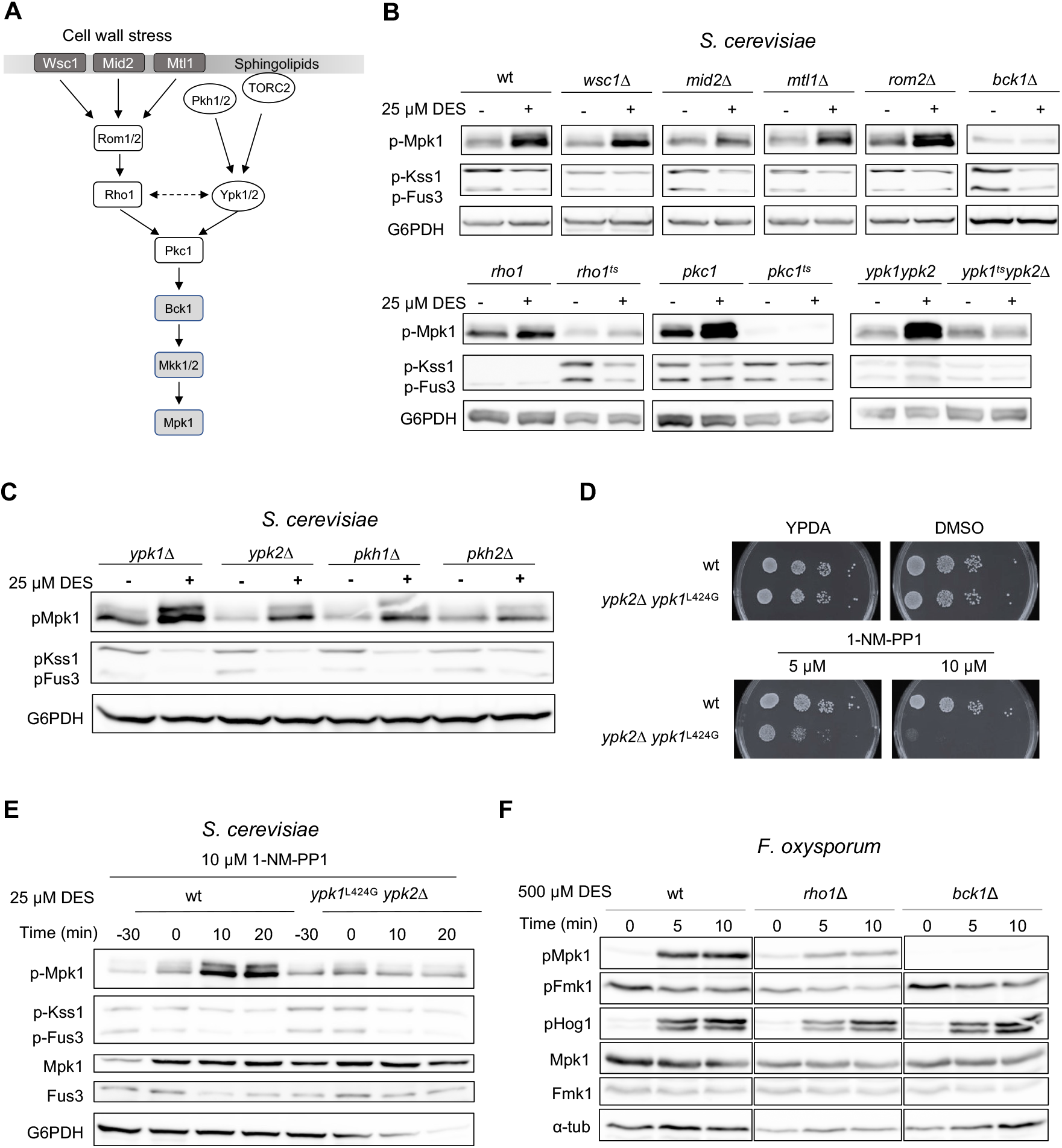
Acid pH-triggered activation of the CWI MAPK cascade is mediated by the Ypk1 sphingolipid signalling branch. **A)** Schematic diagram of the CWI MAPK signaling pathway in *S. cerevisiae.* Modified from (Niles and Powers, 2014). **B,D,E)** Immunoblots showing MAPK phosphorylation in the wild type and in the indicated deletion or temperature-sensitive (ts) mutants of *S. cerevisiae*, at 0 (−) and 20 min (+) (B,D) or at the indicated times (E) after addition of 25 μM DES. In (B), the strains were shifted to the restrictive temperature (34°C) for 60 min before DES addition. In (E), 10 μM of the cell-permeable PP1 analog 1-NM-PP1 was added to the medium 30 min before DES addition. Total protein extracts were subjected to immunoblot with different antibodies as indicated in Fig. 2. **C)** Serial dilutions of the indicated *S. cerevisiae* strains were spotted on YPDA medium supplemented or not with DMSO (solvent) or with the indicated concentrations of the specific Ypk1-AS inhibitor 1-NM-PP1. **F)** Immunoblot showing MAPK phosphorylation in response to 500 μM DES in the wild type and in the indicated mutant strains of *F. oxysporum.* Protein extracts collected at the indicated time points were subjected to immunoblot with different antibodies as indicated in Fig. 2.

Similar to yeast, a partial Rho1 loss-of-function mutant of *F. oxysporum* (Martinez-Rocha *et al.*, 2008) was largely unaffected in DES-triggered phosphorylation of Mpk1 and Hog1 while a mutant lacking the MAPKKK Bck1 exhibited constitutively low levels of Mpk1 phosphorylation, but still showed rapid phosphorylation of Hog1 and dephosphorylation of Fmk1 (**Fig. 6F**). Multiple attempts to obtain deletion mutants in the single *F. oxysporum ypk1* ortholog were unsuccessful suggesting that, as in yeast, Ypk1 is essential in this fungus (**Fig. 6—figure supplement 1A-D**). We therefore recreated the analog-sensitive (AS) *ypk1^L424G^* allele previously used in *S. cerevisiae* (Berchtold *et al.*, 2012) by changing the conserved leucine residue of the *F. oxysporum* Ypk1 to glycine (*ypk1*^L368G^) (**Fig. 6—figure supplement 2A**). A transformant showing homologous replacement of *ypk1* with *ypk1*^L368G^ was confirmed by Sanger sequencing (**Fig. 6—figure supplement 2B**). However, two independent monoconidial isolates of this transformant were insensitive to 1-NM-PP1 and unaffected in DES-triggered Mpk1 phosphorylation (**Fig. 6—figure supplement 2C,D**). We also attempted to recreate the *S. cerevisiae ypk1*-ts allele (Casamayor *et al.*, 1999) by changing the two conserved I^428^ and Y^480^ residues of *F. oxysporum* Ypk1 to T and C, respectively (**Fig. 6—figure supplement 2E**). However, in contrast to yeast, growth of the *F. oxysporum* transformant carrying a homologous insertion of the *ypk1^I428T,Y480C^* allele was not significantly inhibited at high temperature, suggesting that these mutations do not confer temperature sensitivity in this species (**Fig. 6—figure supplement 2F,G**). Collectively, our results suggest that the TORC2-Pkh-Ypk1 upstream module mediates activation of Mpk1 in response to pH_c_ acidification in *S. cerevisiae*, although this role remains to be functionally confirmed in *F. oxysporum*.

**Figure 6—figure supplement 1.**
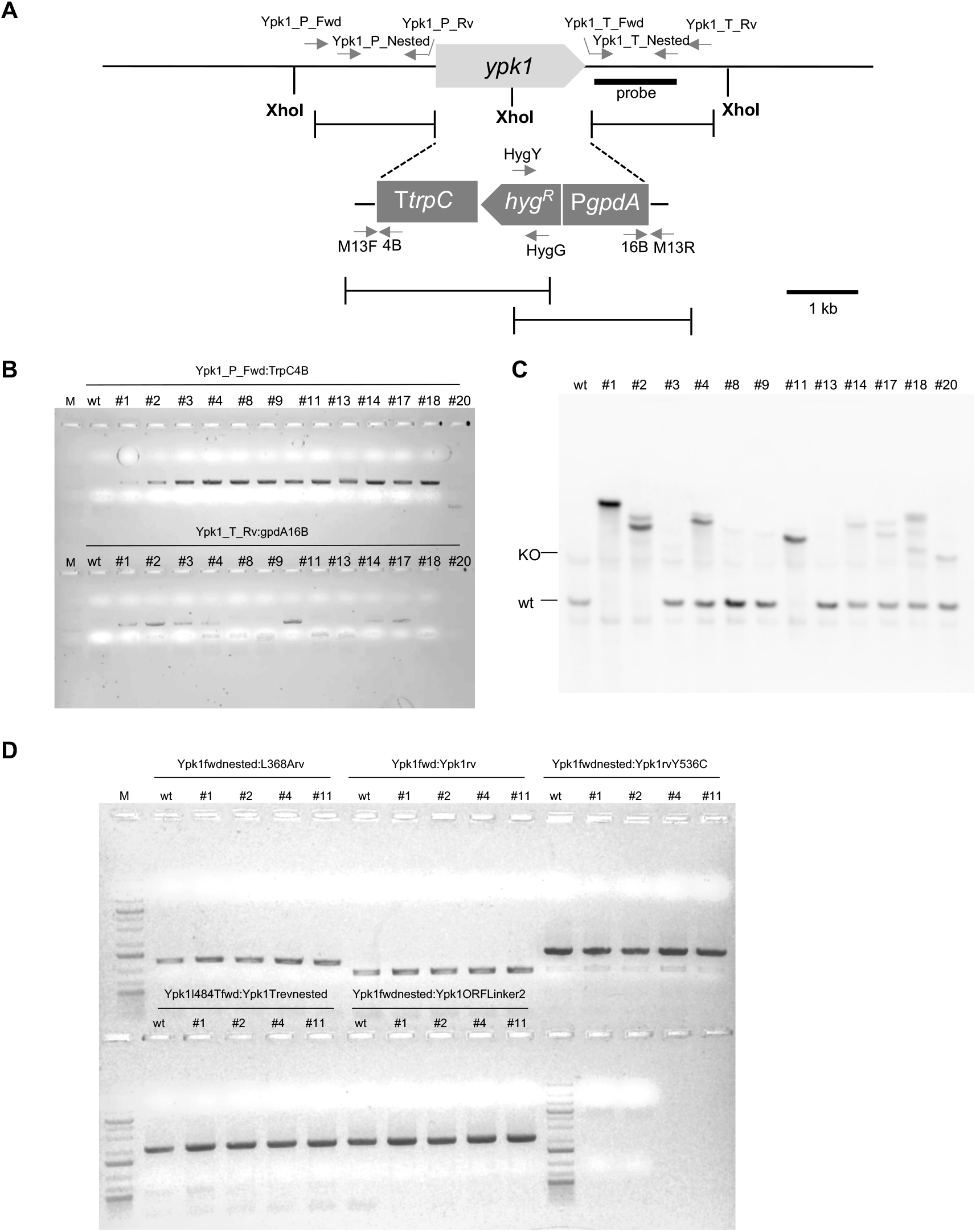
Failure to obtain *ypk1*Δ knockout mutants suggests that Ypk1 is essential in *F. oxysporum.* **A)** Schematic diagram showing the targeted deletion of the *F. oxysporum ypk1* gene using the split-marker method. Gene knockout constructs were obtained by fusion PCR. Relative positions of restriction sites and Southern probes as well as of the PCR primers used are indicated. *hygR*, hygromycin resistance gene; *PgpdA, gpdA* promoter; *TtrpC, trpC* terminator (both from *A. nidulans).* **B,C)** Genomic DNA of the wild type (wt) and independent hygromycin resistant transformants was subjected to PCR with the indicated pairs of primers (B) or treated with *Xho*I (C). The samples were separated on 0.7% agarose gels and imaged (B) or transferred to nylon membranes and hybridized with the DIG labelled DNA probe (C). Relative positions of the expected wild type or knockout (KO) hybridizing bands in (C) are indicated on the left. **D)** Genomic DNA of the wild type (wt) and independent hygromycin resistant transformants was subjected to PCR with the indicated pairs of primers, separated on 0.7% agarose gels and imaged. M, Molecular size markers.

**Figure 6—figure supplement 2.**
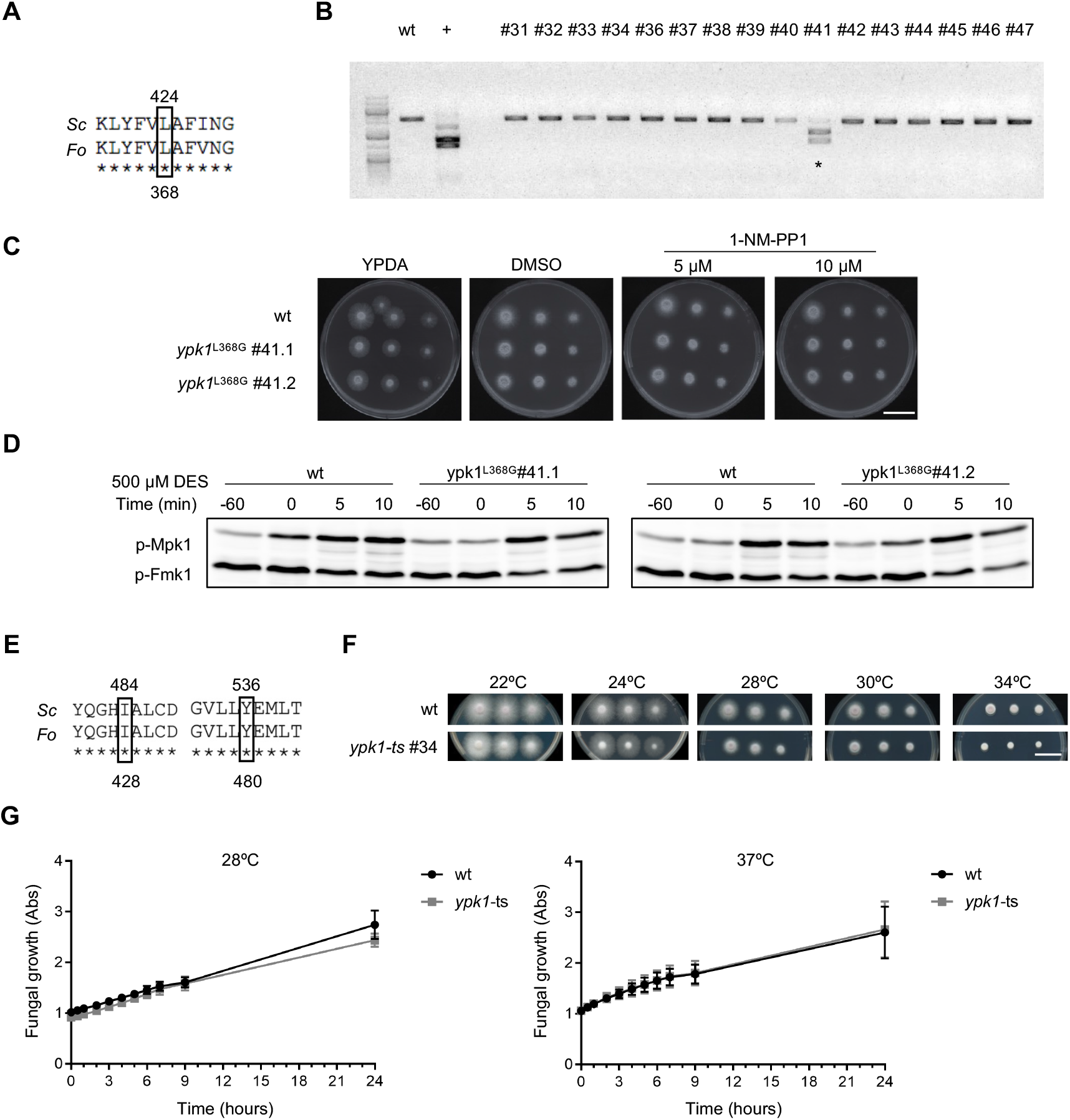
Attempts of mutational analysis of *ypk1* in *F. oxysporum.* **A)** Amino acid alignment showing the contextual conservation in *F. oxysporum* of the L424 residue of *S. cerevisiae* Ypk1, whose mutation to G in yeast causes sensitivity to the analog 1-NM-PP1 (Berchtold *et al.*, 2012). **B)** Screening of *F. oxysporum* transformants for the L368G mutation using RFLP analysis. Genomic DNA of the wild type (wt) and independent hygromycin resistant transformants was subjected to PCR followed by treatment with the restriction enzyme *Nar*I which cuts the DNA fragment carrying the *ypk1*^L368G^ mutation. As a positive control (+), PCR was performed on the DNA construct employed for transformation. **C)** Serial dilutions of fresh microconidia of the wt and two monoconidial isolates of transformant #41 carrying the *ypk1*^L368G^ mutation were spot-inoculated on plates containing YPDA supplemented with the indicated concentrations of 1-NM-PP1. Plates were incubated at 28°C in the dark and imaged after 2 days. Images shown are representative of three independent biologic replicates. Scale bar, 2 cm. **D)** Western blot showing MAPK phosphorylation in response to 500 μM DES in the wt and two monoconidial isolates of transformant #41 carrying the *ypk1*^L368G^ mutation. The specific Ypk1-AS inhibitor 1-NM-PP1 (40 μM) was added 60 min before DES addition (−60). Protein extracts were subjected to immunoblot analysis with anti-phospho-p44/42 MAPK antibody to detect phosphorylated p-Mpk1 and p-Fmk1. **E)** Amino acid alignment showing the contextual conservation in *F. oxysporum* of the 1484 and Y536 residues of *S. cerevisiae* Ypk1, whose simultaneous mutation to T and C, respectively, causes temperature sensitivity in yeast. **F)** Analysis of temperature sensitivity in the *F. oxysporum* wild type strain and *ypk1-ts* #34 transformant carrying the I428T and Y480C mutations. Serial dilutions of fresh microconidia were spot-inoculated on PDA plates, incubated at indicated temperatures in the dark and imaged after 3 days. Images shown are representative of three independent biologic replicates. Scale bar, 2 cm. **G)** Growth of the wt and the y*pk1*-ts #34 strain in PDB at 28°C or 37°C was monitored by measuring absorbance (Abs) at 600 nm. Values were normalized to time zero. Data show the mean ± s.d. of three independent replicates from one representative experiment. Experiments were performed twice with similar results.

### Acidification of pH_c_ triggers changes in long chain base membrane sphingolipids in *F. oxysporum* which are relevant for Mpk1 activation and chemotropism

In *S. cerevisiae*, the activity of Ypk1/2 is regulated by changes in sphingolipid composition of the plasma membrane (Berchtold *et al.*, 2012; García-Marqués *et al.*, 2016; Roelants *et al.*, 2011). Here we found that a downshift of ambient pH or addition of DES in *F. oxysporum* resulted in an increase of the long chain base (LCB) sphingolipid dihydrosphingosine (dhSph) (**Fig. 7A, Fig. 7—figure supplement 1**). A similar increase was also observed in the *mpk1*Δ mutant, suggesting that the LCB response is either independent or upstream of Mpk1. In line with the latter hypothesis, external application of dhSph triggered a rapid phosphorylation response of Mpk1 without affecting the phosphorylation state of Fmk1 (**Fig. 7C**). Importantly, addition of dhSph had no effect on pH_c_ (**Fig. 7B**). These results suggest that Mpk1 activation in response to a downshift of pH_c_ is mediated, at least in part, by an acid-triggered increase in dhSph content.

**Figure 7.**
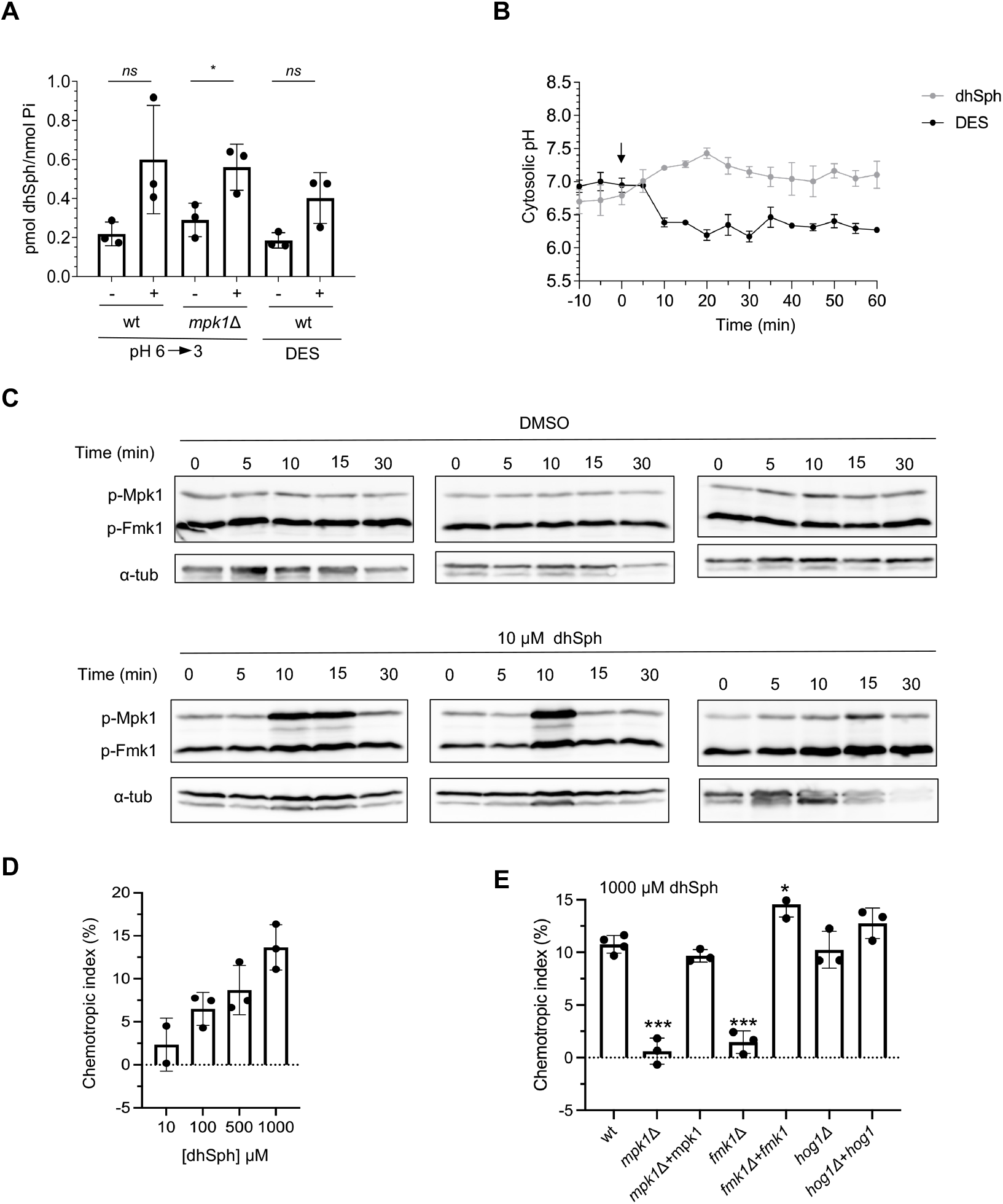
Dihydrosphingosine (dhSph) signals downstream of pH_cy_t to regulate CWI MAPK signalling and hyphal chemotropism. **A)** Acidification of ambient or cytosolic pH leads to increased levels of dhSph. *F. oxysporum* microconidia were pretreated as described in Fig. 2 before shifting the pH of the medium from 6 to 3 with diluted HCl or adding 500 μM DES. Samples were collected before (−) or 10 minutes (+) after the treatment. Extracted lipids were analyzed by HPLC/MS-MS and the dhSph concentration was normalized to phosphate levels (Pi). Data show the mean ± s.d of three independent biological experiments. * p<0.05 *versus* non-treated sample according to Welch’s t-test. **B)** Addition of dhSph does not affect pH_c_yt. *F. oxysporum* microconidia were pretreated as described in Fig. 2 before adding either 500 μM DES or 100 μM dhSph to the medium. pH_c_yt was monitored spectrofluorometrically starting 10 minutes before the treatment. Data show the mean ± s.d. of three independent replicate microwells from one representative experiment. Experiments were performed twice with similar results. **C)** Western blot showing MAPK phosphorylation of *F. oxysporum* in response to addition of the solvent DMSO (upper panels) or 10 μM dhSph (lower panels). Total protein extracts collected at the indicated times were subjected to immunoblot analysis with anti-phospho-p44/42 MAPK to detect phosphorylated p-Mpk1 and p-Fmk1. Anti-α-tubulin (α-tub) was used as loading control. Immunmoblots from 3 independent biological experiments are shown. **D,E)** Directed growth of germ tubes of the *F. oxysporum* wild type strain (D,E) or the indicated mutant strains (E) was determined after 8 h exposure to a gradient of the indicated concentrations of dhSph. *** p<0.001; * p<0.05 *versus* wt according to Welch’s t-test. Data show mean ± s.d of three independent biological experiments (n=500 germ tubes per experiment).

The CWI MAPK cascade was previously shown to mediate chemotropism of *F. oxysporum* towards tomato roots (Turrà *et al.*, 2015). Moreover, it was found that Mpk1 phosphorylation is triggered by plant chemoattractant signals (Nordzieke *et al.*, 2019). Here we found that dhSph acts as a chemoattractant for *F. oxysporum* hyphae. The chemotropic response to dhSph was dose-dependent and required both the Mpk1 and Fmk1 MAPKs (**Fig. 7D,E**). Collectively, these results suggest that dhSph acts as a signal downstream of cytosolic acidification to activate the CWI MAPK signaling cascade and induce hyphal chemotropism.

**Figure 7—figure supplement 1.**
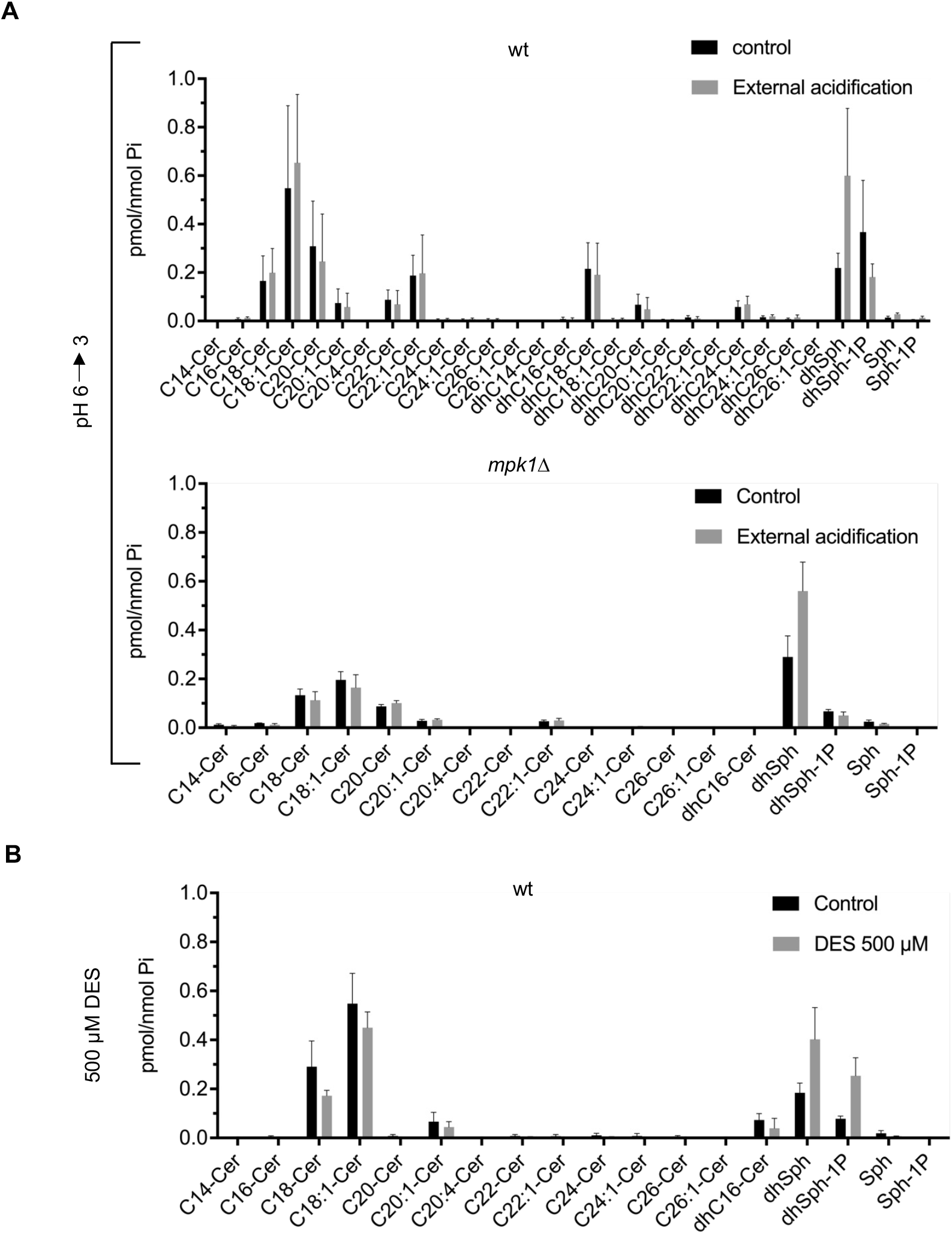
Effect of extra- and intracellular acidification on sphingolipid composition in *F. oxysporum*. **A,B)** Microconidia of the indicated *F. oxysporum* strains were pretreated as described in Fig. 3 before shifting the pH of the medium from 6 to 3 (A) or adding 500 μM DES (B). Samples were collected before (control) or 10 minutes after treatment. Extracted lipids were analyzed by HPLC/MS-MS and the concentration of each ceramide molecular species was normalized to total phosphate levels (Pi). Each graph shows the mean ± s.d. of three independent experiments.

## Discussion

Ambient pH has long been known to act as a key regulator of growth, development and virulence in fungi, but the mechanisms underlying pH control have remained elusive (Fernandes *et al.*, 2017; Peñalva *et al.*, 2014; Vylkova, 2017). We previously found that extracellular alkalinization promotes invasive hyphal growth and plant infection in *F. oxysporum* (Masachis *et al.*, 2016) and that this effect can be reversed by acidification of the rhizosphere, such as that produced by the bacterium *R. aquatilis* through secretion of gluconic acid (Palmieri et *al.*, 2020). Here we show that pH also controls hyphal chemotropism, another infection-related process (Turrà *et al.*, 2015). When exposed to a pH gradient, *F. oxysporum* hyphae displayed robust tropism towards acid. Interestingly, pH regulation of both virulence-related processes is mediated by conserved MAPK cascades. While alkaline-triggered invasive growth functions via activation of the MAPK Fmk1 (Di Pietro *et al.*, 2001; Masachis *et al.*, 2016), chemotropism towards acid is mediated by the CWI MAPK Mpk1 since a downshift of pH induces Mpk1 phosphorylation and loss of Mpk1 impairs acid-triggered tropism. Taken together, these findings highlight the finely tuned functions of different fungal MAPK cascades during infection-related development of *F. oxysporum*, as previously reported, for instance, during appressorial differentiation in the rice blast pathogen *Magnaporthe oryzae* (Cruz-Mireles *et al.*, 2021).

Chemotaxis across pH gradients was previously reported in a variety of organisms. Similar to *F. oxysporum* germ tubes, zoospores of the oomycete pathogen *Phytophthora palmivora* are attracted towards acidic pH (Morris *et al.*, 1995). By contrast, African trypanosomes are attracted to alkali and this response required the cAMP/protein kinase A (PKA) pathway (Shaw *et al.*, 2022). An extracellular pH gradient was shown to act as the dominant cue for the directional migration of MDA-MB-231 tumor cells during hematogenous metastasis (Takahashi *et al.*, 2020). While in eukaryotes the mechanism of chemosensing of pH gradients is largely unknown, it has been elucidated in several bacterial systems. *Escherichia coli* exhibits bidirectional pH chemotaxis allowing it to avoid extreme low and high pH environments, and this behavior is mediated by adaptive methylation of two major chemoreceptors (Yang and Sourjik, 2012). In *Helicobacter pylori*, the causal agent of stomach ulcer, repulsion by acid and attraction towards alkali is important for virulence and requires at least two independent receptors capable of detecting acid gradients (Croxen *et al.*, 2006; Huang *et al.*, 2017). Further work is required to unravel the mechanisms underlying Mpk1-dependent pH tropism of *F. oxysporum.*

### pH_c_ is a signal for regulation of MAPK activity and its downstream responses

To understand how changes in ambient pH cause reprogramming of MAPK phosphorylation, we monitored pH_c_ in a *F. oxysporum* strain expressing the ratiometric pH probe pHluorin. Using two independent methods, spectrofluorometry and confocal microscopy imaging, we determined a homeostatic pH_c_ of 7.3 in *F. oxysporum* hyphae. This value is similar to that reported in *Aspergillus niger* (7.4 to 7.7) using either pHluorin (Huang *et al.*, 2017) or ^31^P-NMR (Hesse *et al.*, 2000). Interestingly, the homeostatic pH_c_ of these two filamentous species is approximately one pH unit higher than that of *S. cerevisiae*, which is around 6.5 (Isom *et al.*, 2013). Here we independently confirmed the difference of approximately one unit between the homeostatic pH_c_ values of *F. oxysporum* and *S. cerevisiae.* Whether the pH_c_ difference is related to the filamentous growth pattern of *Fusarium* and *Aspergillus* as compared to the unicellular lifestyle of budding yeast remains to be determined. Alternatively, the low pH_c_ of *S. cerevisiae* could represent a specific adaptation to its peculiar ecological niche. To test these hypotheses, pH_c_ values should be compared across a broad range of filamentous *versus* non-filamentous fungal species. In the non-filamentous pathogenic yeast *Candida glabrata*, the pH_c_ was around 7.0, which lies between the values of *Fusarium/Aspergillus* and *S. cerevisiae* (Ullah *et al.*, 2013). Interestingly, studies in *Candida albicans* based on laser microspectrofluorimetry or ^31^P-NMR suggested the presence of a pH_c_ gradient along the germ tubes (Rabaste *et al.*, 1995; Stewart *et al.*, 1988), while no evidence for a pH_c_ gradient was found along hyphal tips of pHluorin-expressing *A. niger* hyphae (Bagar *et al.*, 2009).

In *F. oxysporum*, pH_c_ homeostasis responded robustly to shifts in ambient pH. Extreme up- or downshifts of external pH in the range between pH 2 and 9 led to rapid and transient fluctuations in pH_c_, which returned to the homeostatic value within 30 to 60 min. The amplitude of these pH_c_ fluctuations is remarkable, with down- and upshifts of of 1.0 and 0.5 pH units, respectively. A similar range of pH_c_ fluctuations in response to ambient pH shifts was reported in *A. niger* (Bagar *et al.*, 2009). Because of the tight control of pH_c_ in all organisms, even relatively small fluctuations can trigger dramatic cellular responses. For instance in human cells the homeostatic pH_c_ is generally maintained at 7.2, while it is only 0.3-0.5 pH units higher in transformed cells and only 0.3-0.4 units lower in cells triggering apoptosis (Srivastava *et al.*, 2007). Similarly, programmed cell death (PCD) in yeast induced by the antimicrobial protein lactoferrin was preceded by a transient acidification of pH_c_ by 0.3 pH units, whose inhibition prevented PCD indicating that the pH_c_ downshift acts as a triggering signal (Andrés *et al.*, 2019).

Both in *F. oxysporum* and *S. cerevisiae*, extracellular acidification leads to a downshift of pH_c_ and rapid phosphorylation of the two stress-responsive MAPKs Mpk1/Slt2 and Hog1 concomitant with dephosphorylation of the invasive growth MAPK Fmk1/Kss1 and the pheromone response MAPK Fus3, respectively. Crosstalk between the stress and the pheromone response MAPK pathways has been reported in budding yeast, although the underlying molecular mechanisms are poorly understood (Van Drogen *et al.*, 2020). In a previous study, we found that extracellular alkalinization promotes invasive growth and virulence of *F. oxysporum* by triggering rapid phosphorylation of the MAPK Fmk1 (Masachis *et al.*, 2016). Interestingly, pseudofilamentous growth in *S. cerevisiae*, which is mechanistically related to invasive hyphal growth and requires the Fmk1 ortholog Kss1, is also regulated by changes in pH_c_ (Brito *et al.*, 2020). This suggests a conserved role of pH_c_-mediated MAPK regulation in hyphal invasion of the underlying substrate.

Importantly, reprogramming of MAPK phosphorylation in *F. oxysporum* and *S. cerevisiae* was recapitulated in the absence of external pH changes by pharmacological inhibition of the H^+^-ATPase Pma1, which leads to a decrease of pH_c_. These results establish pH_c_ as a key regulator of fungal MAPK activity. Our findings are in line with a previous study in *S. cerevisiae* showing that pH_c_ acts as a second messenger upstream of protein kinase A to regulate the metabolic switch between phospholipid metabolism and lipid storage (Teixeira *et al.*, 2021). Moreover, a rapid downshift of pH_c_ in response to acid stress was proposed to promote cell survival by triggering growth arrest (Lucena *et al.*, 2020). Other stresses such as heat shock or cell wall stress also cause transient intracellular acidification which is associated with increased stress resistance (García *et al.*, 2017; Triandafillou *et al.*, 2020). In animal cells, fluctuations in pH_c_ have been linked to developmental transitions and signaling responses. For instance, intracellular acidification by approximately one pH unit was reported during recovery of developmentally arrested dauer larvae in *Caenorhabditis elegans* (Wadsworth and Riddle, 1988) and in human hippocampal neurons, stimulation with *N-* methyl-D-aspartate (NMDA) causes intracellular acidification (Rathje *et al.*, 2013).

Our finding that inhibition of Pma1 causes a rapid downshift of pH_c_ as well as rapid reprogramming of MAPK phosphorylation confirms the pivotal role of this conserved plasma membrane H^+^-ATPase in pH_c_ homeostasis, cell signaling and development (Kane, 2016). Omeprazole-mediated inhibition of Pma1 in *S. cerevisiae* caused a decrease in pH_c_ and activation of the AMP-activated protein kinase Snf1, suggesting a central role for pH_c_ in the regulation of the cell metabolic program (Salsaa *et al.*, 2021). Intriguingly, unequal distribution of Pma1 between mother and daughter cells was shown to cause pH_c_ asymmetry, with Pma1 accumulating in the aging mother cell while being largely absent from the nascent daughter cell (Henderson *et al.*, 2014). In the human pathogen *C. albicans*, expression of a truncated version of Pma1 led to altered cation responses, disrupted vacuolar morphology and reduced filamentation (Rane *et al.*, 2019). Meanwhile, in the citrus pathogen *Penicillum digitatum* RNAi-mediated silencing of the *pma1* gene caused reduced cell growth and pathogenicity as well as cell wall alterations (Li *et al.*, 2022). Collectively, these results support the role of pH_c_ as a homeostatic sensor that controls the balance between cell growth and stress tolerance (Lucena *et al.*, 2020).

### LCB sphingolipid signaling links pH_c_ acidification withYpk1-mediated activation of the CWI MAPK cascade

By analyzing a battery of *S. cerevisiae* mutants for defects in DES-triggered Mpk1 activation, we identified the AGC kinases Ypk1/2 as key upstream components of the pH_c_-triggered MAPK response. Interestingly, Ypk1/2 was previously detected in a genetic screen for *S. cerevisiae* mutants with increased sensitivity to acetic acid (Mira *et al.*, 2010). Moreover, acetic acid-induced Ypk1 phosphorylation via TORC2 was shown to contribute to cell survival (Guerreiro *et al.*, 2016).

Our attempts to create targeted *ypk1D* knockout mutants in *F. oxysporum* were unsuccessful. This suggests that Ypk1 is essential in *F. oxysporum*, at least under the culture conditions used in this study, and is in line with that reported in *S. cerevisiae* where simultaneous deletion of the paralogs Ypk1 and Ypk2 is lethal. By contrast, in *A. nidulans* and *A. fumigatus*, gene knockout mutants in the single *ypk1* homolog were viable although they showed a drastically sick phenotype and a complete lack of conidiation which precluded their further maintenance and analysis (Colabardini *et al.*, 2013; Fabri *et al.*, 2019). Interestingly, a conditional *Aspergillus* mutant carrying a *ypk1* allele driven by a glucose-repressible promoter exhibited vegetative growth defects, impaired germination and thermosensitivity as well as a decrease in glycosphingolipid (GSL) levels when grown under repressive conditions (Colabardini *et al.*, 2013; Fabri *et al.*, 2019).

In eukaryotic cells, the content of LCBs is generally much lower than that of complex sphingolipids and ceramides and their quantitative balance is tightly regulated. *S. cerevisiae* has two types of LCBs, phytosphingosine and dhSph (Tani, 2016). Here we found that a decrease of pH_c_ in *F. oxysporum*, induced either by extracellular acidification or by pharmacological inhibition of Pma1, led to an approximately two-fold increase of the LCB dhSph. This increase was also observed in a *mpk1D* mutant suggesting that it is independent of Mpk1. Together with the finding that exogenous addition of dhSph causes rapid and transient phosphorylation of Mpk1, our results suggest that dhSph accumulation in response to pH_c_ acidification acts as an activating signal upstream of Ypk1 and the CWI MAPK cascade (**Fig. 8**). In *S. cerevisiae*, LCBs were shown to directly activate Pkh1 and Pkh2, leading to upregulation of Ypk1 and Ypk2 (Liu *et al.*, 2005), while Ypk1 was found to connect sphingolipid biosynthesis with the CWI MAPK pathway (Roelants *et al.*, 2002; Schmelzle *et al.*, 2002). In turn, TORC2-mediated Ypk1 phosphorylation was shown to regulate sphingolipid biosynthesis in response to acetic acid stress (Guerreiro *et al.*, 2016). Lipidomic profiling revealed dramatic changes in sphingolipid composition of *S. cerevisiae* in response to acetic acid stress (Lindberg *et al.*, 2013). Furthermore, LCB levels were transiently increased during heat stress causing a G0/G1 arrest that was essential for thermotolerance (Jenkins *et al.*, 1997).

**Figure 8.**
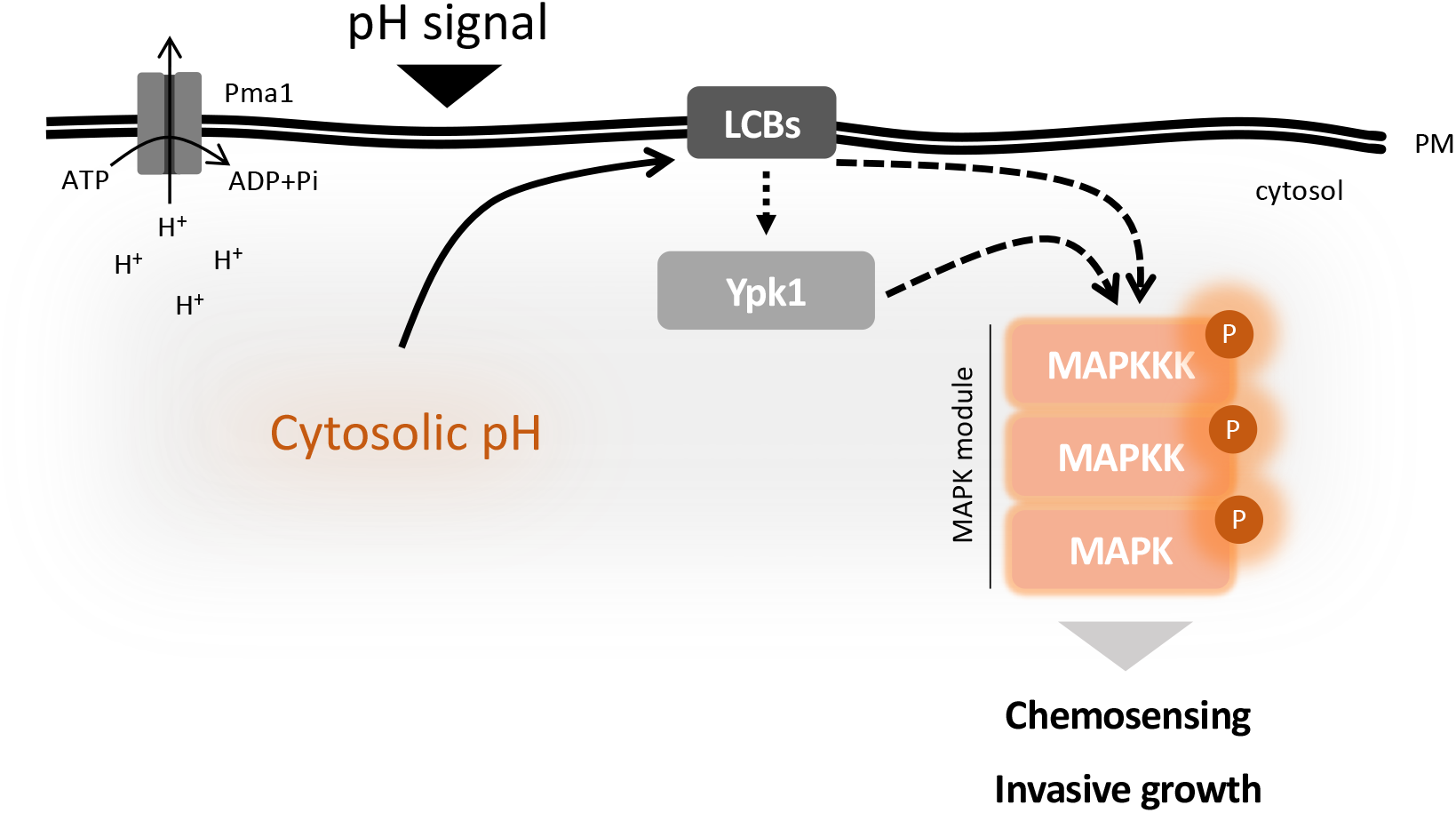
Cytosolic pH controls fungal MAPK signaling and pathogenicity-related functions. The plasma membrane H+-ATPase acts as a key regulator of pH_c_. Acidification of ambient or pH_c_ leads to increased levels of the membrane LCB dhSph which in turn triggers activation of the CWI MAPK Mpk1 via the AGC kinase Ypk1. Acidification of pH_c_ also leads to inactivation of the invasive growth MAPK Fmk1 through an independent mechanism. The interplay of Mpk1 and Fmk1 regulates hyphal chemotropism, invasive growth and virulence of *F. oxysporum.*

Besides the acid stress response, other pH-regulated processes in *F. oxysporum* such as chemotropism of hyphae towards an acid pH gradient could also be regulated by pH_c_-driven fluctuations in LCBs and downstream modulation of Ypk1/Mpk1 activity. Indeed, we found that exogenously applied dhSph acts as a chemoattractant for F. oxysporum hyphae. Interestingly, TORC2-Ypk1 was shown to regulate actin polarization in *S. cerevisiae* by controlling reactive oxygen species (ROS) accumulation via the Pkc1-Mpk1 MAPK cascade (Niles and Powers, 2014). Similarly, mTORC2 plays a conserved role in regulation of the actin cytoskeleton to mediate migration of neutrophils toward chemoattractants (He *et al.*, 2013).

A key finding of our work is that shifts in ambient pH induce dramatic changes in phosphorylation of the three MAPKs of *F. oxysporum*. Using pharmacological inhibition of Pma1, we demonstrate that the MAPK response to acidification is mediated by fluctuations in pH_c_ and LCB content. We speculate that changes in pH_c_ and LCB content may also regulate alkalinization-induced MAPK responses. Intriguingly, the conserved PacC/Rim101 pathway which regulates fungal adaptation to alkaline ambient pH, is also activated by alterations in lipid asymmetry of the plasma membrane (Ikeda *et al.*, 2008). The most upstream component of this pathway, the transmembrane sensor protein PalH/Rim21, was suggested to sense external alkalization through alterations in lipid asymmetry of the plasma membrane via interaction with the inner membrane leaflet (Nishino *et al.*, 2015; Obara and Kamura, 2021). Changes in membrane lipid balance could thus play a key role in sensing of both down- and upshifts of pH_c_. Furthermore, cell signaling by pH_c_ could be mediated by additional mechanisms. For example, cytoplasmic acidification was shown to induce a transition to a solid-like state, which was required for cell survival under conditions of starvation (Munder *et al.*, 2016). Further investigations are needed to fully understand the role of pH_c_ in fungal MAPK signaling, development and virulence.

## Materials and Methods

### Fungal strains and culture conditions

The tomato pathogenic isolate *F. oxysporum* f. sp. *lycopersici* Fol4287 (FGSC 9935) used throughout this study and the strains derived thereof are listed in Supplementary Table S1. Standard methods for fungal growth, handling and genetic transformation were used as described previously (Di Pietro *et al.*, 2001; López-Berges *et al.*, 2012). For measurements of pH_c_ or H^+^-ATPase activity, sphingolipid profiling and western blot analysis, freshly obtained *F. oxysporum* microconidia (5 x 10^6^ spores/ml) were germinated for 15 hours at 28°C and 170 rpm in yeast extract-dextrose (YD) buffered at pH 7.4 with 20 mM HEPES (4-(2-hydroxyethyl)-1-piperazineethanesulfonic acid) unless indicated otherwise. Germlings were washed with KSU buffer (50 mM K_2_HPO_4_, 50 mM sodium succinate, 25 mM urea) adjusted to pH 6.0 with 1.2 M HCl and incubated for 60 min in the same buffer before initiating the experiment. External pH shifts were performed by adding diluted HCl or NaOH. For cell survival assays, samples of 5 x 10^7^ germlings were collected at the indicated time points, serially diluted, spotted on PDA plates, and colony forming units were counted after 48 h incubation at 28°C.

The *S. cerevisiae* wild type and mutant strains used in this study are listed in Supplementary Table S2. Yeast cells were grown overnight in YPD medium at 30°C (25°C for temperature-sensitive strains) at 200 rpm, until reaching the exponential phase (OD64¤nm 0.9). Cells were then transferred to KSU buffer adjusted to pH 6.5 with 1.2 M HCl and incubated for 60 min at 30°C and 200 rpm before initiating the experiment. External pH shifts were performed by adding diluted HCl or NaOH. Temperature-sensitive strains were transferred to the restrictive temperature (34°C) as described (Ozaki *et al.*, 1996).

### Western blot analysis of MAPK phosphorylation

For western blot analysis of MAPK phosphorylation, fungal samples were collected before (time 0) and at the indicated time points after the treatment. *F. oxysporum* samples containing 5 x 10^7^ germlings were collected at each time point and total protein was extracted in lysis buffer (0.2 M NaOH and 0.2% (v/v) β-mercaptoethanol), followed by precipitation with 7.5% (v/v) trichloroacetic acid (TCA) (Méchin *et al.*, 2007). Protein concentration was measured using the Bio-Rad DC™ Protein assay Kit (Bio-Rad, Alcobendas, Spain), using bovine serum albumin as standard. In addition, Coomassie Blue staining was used for visual protein quantification. In *S. cerevisiae*, 25 mL aliquots of exponentially growing cells were collected from each time-point and rapidly quenched by adding 2.0% (v/v) TCA. Protein extraction was performed as described (Isom *et al.*, 2013) with modifications. Briefly, quenched samples were harvested by centrifugation, resuspended in 1 mL of 10 mM sodium azide, centrifuged again and pellets were flash-frozen in liquid nitrogen and stored at −80°C. Frozen pellets resuspended in 0.5 mL of ice-cold TCA buffer (10 mM Tris pH 8.0, 10% (v/v) TCA, 25 mM NH_4_OAc, 1 mM Na2EDTA) were combined with one half volume of glass beads (0.5 mm diameter; Sigma-Aldrich, Madrid, Spain) and vortexed for 3 consecutive cycles of 90 sec interrupted by 30 sec intervals on ice. Precipitated protein was harvested by centrifugation at 13,400 rpm for 10 minutes at 4°C and resuspended in 75 μL resuspension buffer (100 mM Tris, 3% (v/v) SDS, pH 11.0). Protein concentration was measured using the Bio-Rad DC™ Protein assay Kit (Bio-Rad), using bovine serum albumin as standard.

Western blot analysis for determining MAPK phosphorylation status was performed as previously described (Nordzieke *et al.*, 2019). Briefly, protein samples were separated in 10% SDS-polyacrylamide gels and transferred to a nitrocellulose membrane using a Trans-Blot Turbo RTA Midi Nitrocellulose Transfer Kit (Bio-Rad). Phosphorylation of Mpk1 and Fmk1 MAPKs was detected using rabbit anti-Phospho-p44/42 MAPK (Erk1/2) antibody (Thr202/Tyr204, #4370; Cell Signaling Technology, Danvers, MA). Phosphorylation of Hog1 MAPK was detected using rabbit anti-Phospho-p38 MAPK antibody ((Thr180/Tyr182, #9211; Cell Signaling Technology). Total MAPK proteins were detected using a commercial mouse monoclonal anti-Mpk1 antibody (sc-165979, Santa Cruz Biotechnology, Heidelberg, Germany), a custom-designed polyclonal anti-Fus3/anti-Fmk1 antibody (SICGEN Research and Development in Biotechnology Ltd, Cantanhede, Portugal) based on amino acids 38-59 of the predicted *F. oxysporum* Fmk1 protein, or a commercial rabbit polyclonal anti-Hog1 antibody (sc-79079, Santa Cruz Biotechnology). Mouse anti-α-tubulin antibody (#T9026, Sigma-Aldrich) and rabbit anti-glucose-6-phosphate dehydrogenase (G6PDH) (#A9521, Sigma-Aldrich) were used as loading controls for *F. oxysporum* and *S. cerevisiae*, respectively. Hybridising bands were visualized using the ECL™ Select western blotting detection reagent (GE Healthcare, Chicago, IL, USA) in a LAS-3000 detection system (Fujifilm España, Barcelona, Spain).

### Cytosolic pH measurements using the ratiometric fluorescent probe pHluorin

A *F. oxysporum* strain expressing the pH-sensitive GFP variant pHluorin was obtained by co-transformation of fungal protoplasts with the hygromycin resistance cassette (*Hyg*^R^) (Punt *et al.*, 1987) and a PCR fusion construct containing the *pHluorin* gene and the *S. cerevisiae adh5* terminator (Miesenböck *et al.*, 1998) fused to the strong constitutive *Aspergillus nidulans gpdA* promoter (Punt *et al.*, 1987). Hygromycin resistant transformants were screened for the presence of the pHluorin expression cassette using PCR amplification with primers gpda15b and pHLNestRev and confirmed by fluorescence microscopy. Among the obtained transformants carrying the pHluorin cassette, the strain displaying the strongest intracellular fluorescence was selected for further studies. Measurements of pH_c_ in *F. oxysporum* were performed as described (Fernandes *et al.*, 2022), either spectrophluorometrically in microtiter wells or by fluorescence confocal microscopy of single germlings. For spectrophluorometric measurements of pH_c_, germinated microconidia were transferred to KSU at pH 6.0 in 96-well microtiter plates and incubated for 30 minutes at 28°C before reading fluorescence intensities. Fluorescence emission at 510 nm after excitation at 395 nm and 475 nm was monitored over time in a TECAN spectrofluorometer (Infinite M200 PRO, TECAN Life Sciences, Switzerland). After subtracting the values of the pHluorin-negative wild type background for each wavelength, the 395/475 nm ratio was calculated and converted to pH_c_ values using a pH calibration curve obtained with nigericin-permeabilized cells (Fernandes *et al.*, 2022). Each experiment represents the average and standard deviation of three independent replicate wells. Experiments were performed at least twice.

For single-cell analysis of pH_c_, fluorescence intensities were recorded as described (Fernandes *et al.*, 2022) using a Zeiss LSM880 laser confocal microscope equipped with diode (405 nm) and Argon (488 nm) lasers, using a Plan Apo 63x oil 1.4 NA objective. Images were set to 8 bits and the background was subtracted using the lookup table HiLo (Image/Fiji). Cell shape was delimited by drawing a line, the fluorescence intensity was measured within the line for each wavelength, and the 405/488 nm ratio was determined and converted to pH_c_ values using a pH calibration curve obtained with nigericin-permeabilized cells. Each experiment represents the average and standard deviation of at least three independent cells measured.

For measuring pH_c_ in *S. cerevisiae*, the wild type strain BY4741 was transformed with the pYEplac181 plasmid (amp, LEU2) containing the *pHluorin* gene under control of the *TEF1* promotor (Isom *et al.*, 2013). Exponentially growing wild type and pHluorin expressing strains were resuspended in KSU pH 6.5, aliquoted into a 96-well microtiter plate and incubated for 30 minutes at 30°C before reading fluorescence intensities. Measurements and calculations of pH_c_ were performed as described above. Each experiment represents the average and standard deviation of three independent replicate wells. Experiments were performed at least twice.

### Determination of Pma1 H^+^-ATPase activity

For determination of the activity of the plasma membrane H^+^-ATPase Pma1 in *F. oxysporum*, plasma membrane fractions were obtained as previously described (Kahm *et al.*, 2012) with minor modifications. Briefly, samples of 1.25 x 10^8^ germlings per time point were rapidly harvested by filtration through a nylon filter (mesh size 10 μm) and flash-frozen in liquid nitrogen. For crude membrane purification, mycelia were resuspended in 3 mL extraction buffer (0.3 M Tris-HCl pH 8.0, 0.3 M KCl, 30 mM EDTA, 5.3 mM dithiothreitol) supplemented with 40 μL protease inhibitor cocktail (Roche Life Sciences, Barcelona, Spain). Then, precooled 0.5 mm glass beads (5 mL per sample) were added and samples were vortexed for 3 consecutive cycles of 90 sec interrupted by 30 sec on ice. Cell lysates were centrifuged for 5 minutes at 1157 g, and the supernatant was further centrifuged for 20 minutes at 18,472 g. Pellets were resuspended in a mixture of 100 μL glycerol buffer (20% (v/v) glycerol, 10 mM Tris-HCl pH 7.6, 1 mM EDTA, 1 mM DTT) and 900 μL of cold ultrapure water and centrifuged 30 minutes at 18,472 g to remove inorganic phosphate and other contaminants. Finally, the membrane fraction was resuspended in 100 μL glycerol buffer and the diethylstilbestrol (DES)-sensitive ATPase activity was measured as decribed (Kahm *et al.*, 2012), with minor changes. Briefly, samples with 6 μg membrane extracts were assayed for ATPase activity in a 96-well microtiter plate in the presence of 0.2 mM of the Pma1-specific inhibitor DES or methanol (solvent control). The plate was incubated for 30 minutes at RT to allow irreversible inhibition of Pma1 activity by DES. Then, ATP-containing buffer (50 mM MES-Tris pH 5.7, 5 mM MgSO_4_, 50 mM KNO_3_, 5 mM sodium azide, 0.3 mM ammonium molybdate, 2 mM ATP) was added and samples were incubated for 40 min at 30°C. The reaction was stopped by adding detection buffer (2% (v/v) sulfuric acid, 0.5% (w/v) ammonium molybdate, 0.5% (w/v) SDS and 0.1% (w/v) ascorbic acid) and incubated for 20 min before reading absorbance at 750 nm in a TECAN spectrofluorometer. Specific Pma1 H^+^-ATPase activity was calculated by subtracting the residual activity value obtained in the presence of DES from the total activity (methanol), expressed in mmol/min/g protein assayed and normalized to time point zero for each time-course. The results represent the average and standard deviation of three independent replicates. Experiments were performed at least twice.

### Quantification of hyphal chemotropism and invasive hyphal growth

Chemotropic growth was measured using a quantitative plate assay described previously (Turrà *et al.*, 2015). Briefly, 10^6^ microconidia were embedded in 0.5% water agar, incubated 8 hours at 28 °C in the presence of a chemoattractant gradient, and the direction of germ tubes relative to a central scoring line was determined in an Olympus binocular microscope at 9,200 x magnification. For each sample, five independent batches of cells (n = 100 cells per batch) were scored. Calculation of the chemotropic index and statistical analysis was done as described (Turrà *et al.*, 2015). Experiments were performed at least three times with similar results. For pH chemotropism, a gradient competition assay (Turrà *et al.*, 2015) was performed between two wells at both sides of the scoring line containing 25mM HCl or NaOH, respectively, as chemoattractants.

Invasive hyphal growth through cellophane membranes was determined as described (López-Berges *et al.*, 2010). Briefly, potato-dextrose-broth (PDB) agar plates buffered to pH 5 or pH 7 with 100 mM MES (2-(N-morpholino)ethanesulfonic acid) were covered with a cellophane membrane and 5 x 10^4^ microconidia were spot-inoculated on the top at the center of the plate. After 2 days of incubation at 28°C, the cellophane membrane with the fungal colony was carefully removed and plates were incubated for 1 additional day at 28°C. Plates were scanned before and after cellophane removal. Triplicates were performed for each strain and condition, and two independent experiments were performed with similar results.

### Sphingolipid profiling

For quantitative analysis of sphingolipid species, samples of 5 x 10^7^ germlings were collected before (time 0) and at the indicated time-points after the treatment. Samples were flash-frozen in liquid nitrogen, lyophilized and submitted to quantitative sphingolipid analysis at the Lipidomics Shared Resource, Medical University of South Carolina, USA. The levels of the ceramide long-chain (sphingoid) bases (LCB) sphingosine (Sph) and dihydrosphingosine (dhSph), as well as sphingoid base-1-phosphates (S1P and dhS1P) and ceramide molecular species were measured by high-performance liquid chromatography/mass spectrometry (HPLC-MS/MS) (Bielawski *et al.*, 2010). Quantitative analysis of sphingolipids was based on eight-point calibration curves generated for each target analyte. Synthetic standards along with a set of internal standards were spiked into an artificial matrix and then subjected to an identical extraction procedure to that of the biological samples. The extracted standards were analyzed by the HPLC/MS-MS operating in positive multiple reaction-monitoring (MRM) mode employing a gradient elution. Analyte-specific calibration curves were generated by plotting the analyte/internal standard peak area ratios against the analyte concentrations. Lipids with no authentic standards were quantitated using the calibration curve of their closest counterpart. The concentration of each sphingolipid species was normalized to the total phosphate level in each biological sample.

## Acknowledgements

We thank Esther Martínez Aguilera and María Ortega Bellido for valuable technical assistance and Profs. Zdena Palková, Charles University, Prague, Czech Republic and Henrik G. Dohlman, University of North Carolina, Chapel Hill, USA, for kindly providing the pHluorin gene and plasmid pYEplac181, respectively. Confocal microscopy was carried out at the Central Service for Research Support (SCAI) of the University of Córdoba. This work was supported by grants from the Spanish Ministry of Science and Innovation (MICINN, PID2019-108045RB-I00) and Junta de Andalucía (P20_00179) to A.D.P and grant PID2019-105342GB-I00/AEI/10.13039/501100011033) to H.M. and T.F-A. T.R.F. was supported by the Marie Curie ITN FUNGIBRAIN (FP7-PEOPLE-ITN-607963). M.M.G. was supported by a FPI predoctoral fellowship from MICINN (BES-2017-082775).

## Author contributions

Tânia R. Fernandes, Conceptualization, Formal analysis, Investigation, Methodology, Resources, Validation, Visualization, Writing - original draft, Writing - review and editing, Incorporating revisions from co-authors; Melani Mariscal, Conceptualization, Formal analysis, Investigation, Methodology, Resources, Validation, Visualization, Writing - review and editing; Antonio Serrano, Conceptualization, Investigation, Methodology, Resources, Validation, Visualization, Writing - review and editing; David Segorbe, Conceptualization, Investigation, Methodology, Resources, Validation; Teresa Fernández-Acero, Methodology, Resources, Validation; Humberto Martín, Conceptualization, Funding acquisition, Investigation, Methodology, Resources, Writing - review and editing; David Turrà, Conceptualization, Investigation, Formal analysis, Methodology, Resources, Writing - review and editing; Antonio Di Pietro, Conceptualization, Visualization, Funding acquisition, Project administration, Supervision, Writing - original draft, Writing - review and editing, Incorporating revisions from co-authors, Writing the final draft

## Additional files

### Supplementary files

- Supplementary Table 1. *Fusarium oxysporum* strains used in this study.
- Supplementary Table 2. *Saccharomyces cerevisiae* strains used in this study.

### Data and materials availability

All data needed to evaluate the conclusions in the paper are present in the paper and/or the Supplementary files.

---

**Table 1.** *Fusarium oxysporum* strains used in this study.

| Strain | Genotype | Reference |
| --- | --- | --- |
| FGSC 4287 | Wild type | (Di Pietro *et al.*, 2001) |
| *bck1Δ* | *bck1::HYG* | (Turrà *et al.*, 2015) |
| *fmk1Δ* | *fmk1::PHLEO* | (Di Pietro *et al.*, 2001) |
| *fmk1Δ hog1Δ* | *fmk1::PHLEO; hog1::HYG* | (Segorbe *et al.*, 2017) |
| *fmk1Δ+fmk1* | *fmk1::PHLEO; fmk1+HYG* | (Di Pietro *et al.*, 2001) |
| *hog1Δ* | *hog1::HYG* | (Segorbe *et al.*, 2017) |
| *hog1Δ+hog1* | *hog1::HYG; hog1+PHLEO* | (Segorbe *et al.*, 2017) |
| *mpk1Δ* | *mpk1::HYG* | (Turrà *et al.*, 2015) |
| *mpk1Δ fmk1Δ* | *mpk1::HYG; fmk1::PHLEO* | (Turrà *et al.*, 2015) |
| *mpk1Δ hog1Δ* | *mpk1::HYG; hog1::PHLEO* | (Segorbe *et al.*, 2017) |
| *mpk1Δ+mpk1* | *mpk1::HYG; mpk1+PHLEO* | (Turrà *et al.*, 2015) |
| *pacCΔ* | *pacC::HYG* | This study |
| *pacCΔ::pHluorin* | *pHluorin; HYG; pacC::NEO* | This study |
| *palHΔ* | *palH::HYG* | This study |
| *palHΔ::pHluorin* | *pHluorin; HYG; palH::NEO* | This study |
| *pHluorin* | *pHluorin; HYG* | (Fernandes *et al.*, 2022) |
| *rho1Δ* | *rho1::HYG* | (Martinez-Rocha *et al.*, 2008) |

**Table 2.** *Saccharomyces cerevisiae* strains used in this study.

| Strain | Gentotype | Reference |
| --- | --- | --- |
| BY4741 | *MATa his3Δ1 leu2Δ0 met15Δ0 ura3Δ0* | Euroscarf |
| pHluorin | BY4741; pYEplac181-Tef1-pHluorin | (Isom *et al.*, 2013) |
| *bck1Δ* | BY4741 *bck1::KanMX4* | Euroscarf |
| *mid2Δ* | BY4741 *mid2::KanMX4* | Euroscarf |
| *mtl1Δ* | BY4741 *mtl1::KanMX4* | Euroscarf |
| *pkh1Δ* | BY4741 *pkh1::KanMX4* | Euroscarf |
| *pkh2Δ* | BY4741 *pkh2::KanMX4* | Euroscarf |
| *rom2Δ* | BY4741 *rom2::KanMX4* | Euroscarf |
| *wsc1Δ* | BY4741 *wsc1::KanMX4* | Euroscarf |
| *ypk1Δ* | BY4741 *ypk1::KanMX4* | Euroscarf |
| *ypk2Δ* | BY4741 *ypk2::KanMX4* | Euroscarf |
| YPH499 | *MATa ade2-10 trp1-63 leu2-1 ura3-52 his3-Δ200 lys2-801* | (Sikorski and Hieter, 1989) |
| *ypk1-ts/ypk2Δ* | YPH499 *ypk1-ts::HIS3 ypk2::TRP1* | (Casamayor *et al.*, 1999) |
| OHNY | *MATa ura3 his3 trp1 leu2 ade2* | (Ozaki *et al.*, 1996) |
| *rho1-ts* | OHNY *rho1-104* | (Ozaki *et al.*, 1996) |
| SEY6221 | *MATa leu2-3,112 ura3-52 his3Δ200 trp1Δ901 suc2Δ9 ade2-101* | (Paravicini *et al.*, 1992) |
| *pkc1-ts* | SEY6221 *pkc1Δ1::HIS3* | (Paravicini *et al.*, 1992) |
| TB50a | *MATa trp1 his3 ura3 leu2* | (Berchtold *et al.*, 2012) |
| *ypk1*<sup>L424G</sup><br>*ypk2Δ* | TB50a *ypk1Δ::KANMX4 ypk2Δ::HIS3* pRS416-*ypk1*<sup>L424G</sup> | (Berchtold *et al.*, 2012) |

